# Making invisible excited state protein structures visible by combining NMR and machine learning

**DOI:** 10.1101/2025.11.27.690854

**Authors:** Jeffrey P. Bonin, Jin Sub Lee, Zi Hao Liu, Philip M. Kim, Lewis E. Kay

## Abstract

NMR relaxation studies of the precursor form of the pro-inflammatory cytokine interleukin-18, pro-IL-18, show that it adopts two sparsely populated (<0.5%) and transiently formed (ms lifetimes) excited state conformations in exchange with a highly populated ground state conformer. Although NMR data localize regions undergoing exchange to a pair of short β-strands that are preserved in at least one of the excited states, additional structural information is not forthcoming. Here we develop a protocol whereby the NMR data is used to select alternative conformers of pro-IL-18 from ensembles predicted by the generative ML model AlphaFlow that are then evaluated through further NMR experiments. The approach identifies distinct excited state conformers and suggests a general method for combining experiment with computation to characterize protein energy landscapes.

## INTRODUCTION

Proteins are dynamic molecules, and their dynamics play critical roles in their function (1). Motions can occur on timescales ranging from femtoseconds (fs) to many seconds (s) or hours (h), often involving significant rearrangements of structure to accommodate specific interactions with targets or to respond to various stimuli in the environment (2). In many cases these dynamics involve exchange between ground (lowest energy, highly populated) and excited (higher energies, sparsely populated) conformational states, the latter of which can play critical roles in molecular recognition, including binding of ligands to proteins as cancer therapeutics (3), signaling (4), enzyme catalysis (5–8), and protein folding and misfolding processes (9–11). Although ground state conformers are characterized in detail using well established biophysical techniques, excited states are sparsely populated and often transiently formed so that they are rendered invisible to many of the most used biophysical methods. While computational approaches are now widely available for predicting structures of ground state conformations (12,13), they are much less efficient in generating accurate representations of excited states for which little experimental data is available for training (14).

Nuclear Magnetic Resonance (NMR) spectroscopy is a powerful tool for studies of protein dynamics over timescales ranging from ps to h. The development of relaxation-based experiments such as Carr-Purcell-Meiboom-Gill (CPMG) relaxation dispersion (15–17) and Chemical Exchange Saturation Transfer (CEST) (18) has enabled, in some cases, atomic resolution descriptions of excited conformational states that cannot be directly observed in NMR spectra, as well as quantification of the kinetics and thermodynamics involved in the underlying conformational exchange process(es). Most applications have involved excited states with significant changes to secondary structural elements relative to the ground state, or with rearrangements of domains, and have focused on conformers that originate due to thermal fluctuations from the ground state (9,10,19–23). Related relaxation experiments, termed Dark State Exchange Saturation Transfer or DEST have been used to study exchange between molecules free in solution and bound to the surface of large macromolecules to obtain structural information on the bound conformation(s) of the ligand and the kinetics and mechanism of the binding process (24). Pressure-based NMR methods offer an exciting route to characterize transient intermediates along reaction trajectories in cases where the states involved differ in volume (25–27).

Despite the underlying potential of the various NMR methods there are limitations, as we recently encountered in CPMG and CEST relaxation studies of pro-IL-18 (28). Pro-IL-18 is activated via cleavage by caspases-1 and −4, removing the first 36 residues to produce the mature form of the protein (29,30). Previously, we showed that pro-IL-18 transiently adopts two excited states (ES_1_ and ES_2_) on the ms timescale, with chemical shift differences between each of them and the ground conformation localized to a region of the protein containing two short β strands, β1 and β* (**Fig. 1**), which are involved in caspase binding. The chemical shift differences are especially large within and adjacent to β* (**Fig. 1**). Backbone chemical shifts indicate that these β strands do not unfold in either excited state and maintain their secondary structures in at least one of the rare conformations for the wild-type (WT) protein. Further structural insights from the NMR data could not be obtained, however, because the shifts themselves provide little information about the orientation of the strands with respect to the remaining structure, and orientational restraints derived from anisotropic interactions such as residual dipolar couplings can be difficult to quantify in exchanging systems with more than two states.

**Figure 1:**
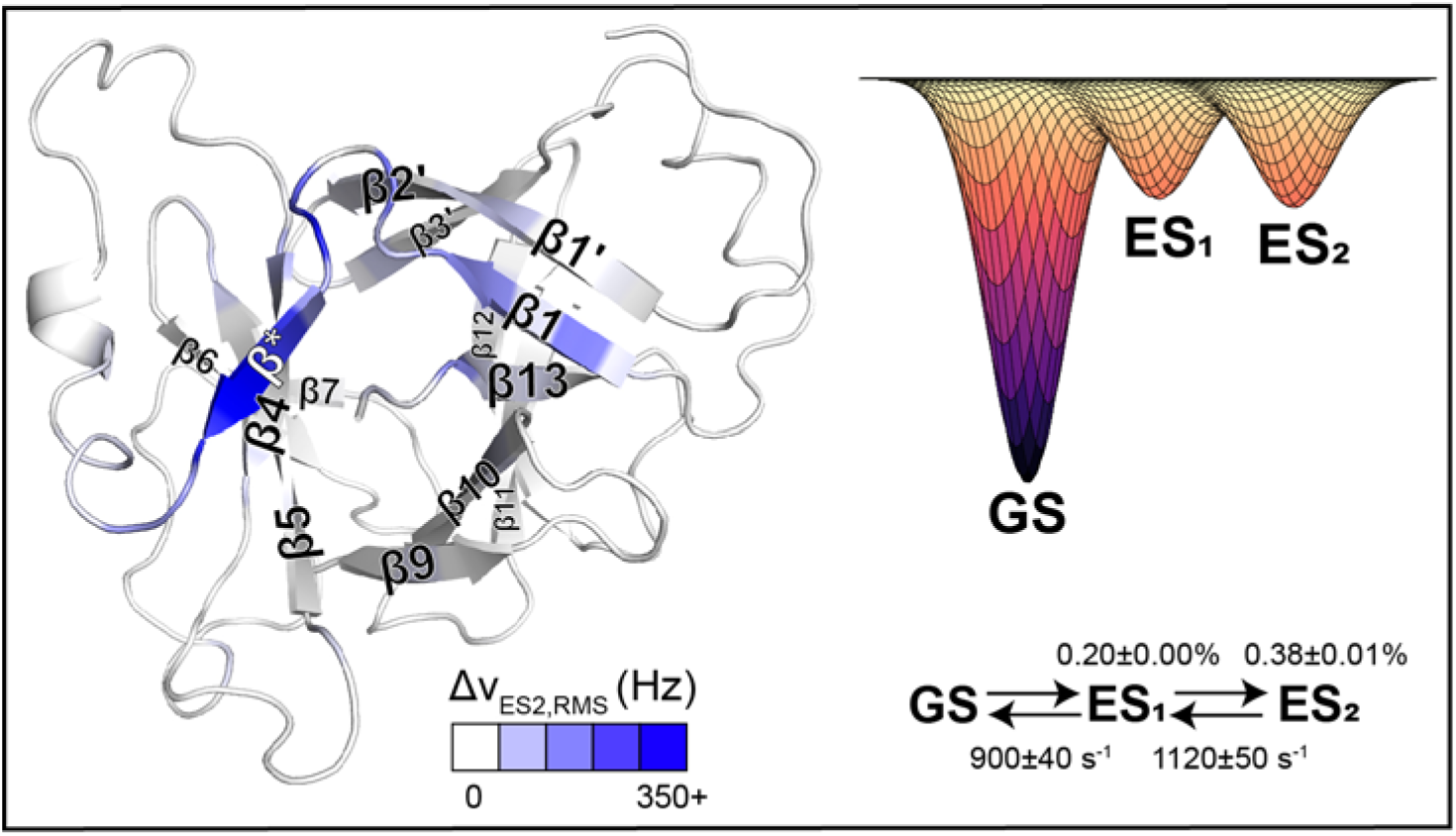
Pro-IL-18 adopts invisible, excited states in regions proximal to the binding interface with caspase-1. Pro-IL-18 accesses two excited states (ES_1_ and ES_2_), with the largest chemical shift changes between ground and excited states occurring in and adjacent to the β* strand (28). The root mean squared difference in chemical shifts between the WT ground state (GS) and ES_2_, Δν_ES2,RMS_ (Hz), calculated based on ^1^HN, ^15^N, ^13^CO, ^13^C^α^, ^13^C^β^, and ^1^H^α^ shifts recorded at a field of 23.4 T (1 GHz), is shown on the structure as a white-blue gradient. The fitted exchange scheme, including rates of interconversion (sums of forward and backwards rates connecting pairs of states) and populations, 25 °C, is shown.

We wondered if new machine learning-based tools might be able to provide some insights into excited state structures in cases where secondary structure is preserved, but where significant shift differences between ground and excited conformers indicate distinct tertiary arrangements, as for pro-IL-18. Recent advances in machine learning allow for rapid generative modeling of protein conformational ensembles (31–33), including for fold-switchers (34–36), but thus far only for conformations which are similarly stable to the ground state. We here thought to identify conformations of higher energy, excited (invisible) states by combining a large ensemble of diverse structures obtained from AlphaFlow, a recently developed generative framework, with knowledge of the pro-IL-18 excited states from initial CPMG and CEST NMR measurements. In this manner, structures within the ensemble were identified that could resemble the excited states. These potential structures were then used to suggest further NMR experiments which validated the AlphaFlow predictions, hence providing atomic-resolution models of the excited state conformers identified through NMR relaxation experiments.

## RESULTS

### AlphaFlow predicts two alternative folded conformations of β*

To gain structural insights into the excited states of pro-IL-18 that can then be used to guide their further experimental characterization, beyond what was available through initial NMR relaxation studies, we turned to AlphaFlow. The amino acid sequence of pro-IL-18 was fed to AlphaFlow, along with its NMR structure which was provided as a template. We found that the template structure was required to prevent AlphaFlow from generating erroneous structures, similar to AlphaFold’s incorrectly predicted, high confidence structure of pro-IL-18 (37) that did not incorporate the pro-sequence properly within the folded domain of the structure. To generate structures with greater variability within the region of interest, the coordinates of residues L45-Q60 (containing β1 and β*) were masked (i.e. withheld) in the template structure. Using this approach, 30,000 structures of pro-IL-18 were produced, showing variability within and adjacent to the β* strand. The ability to mask only the region of interest was key, as was the generation of a large ensemble, to reveal a variety of states including hidden, excited conformers. These requirements favor a rapid generative model such as AlphaFlow. Principal component analysis was used to identify distinct conformations among these structures; however, we found that there were no clear groupings of the structures in the first two principal components (**fig. S1**).

Instead, we utilized our knowledge of the excited state structures from our initial NMR characterization to search for relevant groups of conformers within the AlphaFlow set. Using a procedure which included groupings based on (i) the backbone dihedral angles of Q54 where ψ changes from ∼0° (ground state, GS) to 120° (ES_2_) as predicted by the program TALOS (38), which uses input backbone chemical shifts to generate the associated dihedral angles (see supplementary materials), and (ii) the backbone dihedral angles of the residues forming β* (V55-I58), which are still in the appropriate range for a β strand for at least one of the excited states, as well as additional criteria described in Materials & Methods and illustrated in **fig. S2,** five groups of structures were identified (**Fig. 2A**). These were: (1) a ground state-like group (GS), which is nearly identical to our experimentally determined structure of pro-IL-18; (2) an unfolded group (Unfolded), where residues V55-I58 do not adopt β strand-like backbone dihedral angles; (3) an alternative folded group (Alt1), where the β* strand has flipped around; (4) a second alternative folded group (Alt2), where the β* strand is again flipped, but shifted “down” by two residues relative to Alt1; (5) a final group (Other) where the orientation of β* has rotated by approximately 90° from the GS conformation in many of the structures.

**Figure 2:**
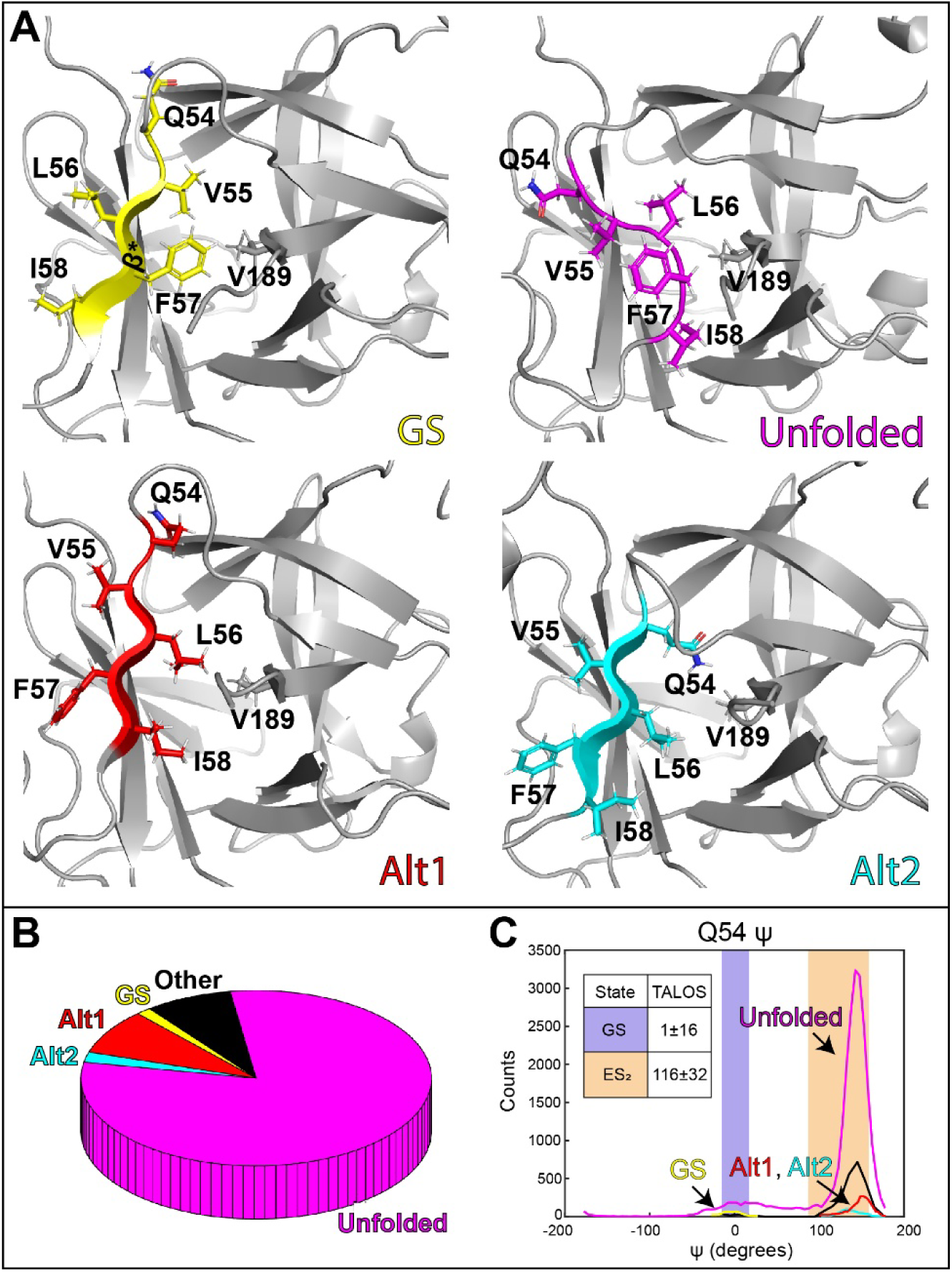
AlphaFlow predicts two alternative folded conformations of β*. (A) Representative structures from four different classes generated by AlphaFlow: GS (similar to the experimentally determined structure), Unfolded (residues V55-I58 are disordered), Alt1 (β* strand is flipped around, shifted “up” by one residue), and Alt2 (β* strand is flipped around, shifted “down” by one residue). (B) Relative abundance of the structural classes across all AlphaFlow structures. (C) Q54 ψ angle distribution from all AlphaFlow structures (*n* = 29,950) is largely bimodal, with one mode centered about ∼0° degrees (corresponding to GS), and a second mode centered on ∼145° (β strand-like, corresponding to the alternative conformations, Alt1 and Alt2, as well as the ‘Unfolded’ and ‘Other’ classes of structures). Notably, the Q54 ψ angle prediction from TALOS for WT ES_2_ (28) (inset) and for ES_1_ and ES_2_ of the Q54V variant (present work, **table S1**) agrees with the ψ angles found in the Alt1 and Alt2 structures.

Most of the structures generated by AlphaFlow belong to the unfolded group, and both the Alt1 and Alt2 groups contain more structures than the GS group (**Fig. 2B**), indicating that AlphaFlow does not accurately capture the relative abundances of various structures within the conformational ensemble of pro-IL-18. However, as AlphaFlow predicts two main alternative structures, Alt1 and Alt2, while our NMR experiments show that pro-IL-18 adopts two excited states, we wondered whether these predicted structures might resemble the sparse conformers identified by NMR. Alt1 and Alt2 are consistent with our initial NMR characterization both in terms of the ψ angles of Q54 for both alternative forms, which agree with the β strand-like prediction of TALOS for the Q54 ψ angle of ES_1_ (**table S1**) and ES_2_ (**Fig. 2C**), and with the secondary structure of V55-I58, a region which forms a β strand in both alternative structures as well as in ES_1_ (**table S1**) and ES_2_ (28).

### Replacing Q54 with a **β** strand-favoring residue increases excited state populations

The backbone dihedral angles of Q54 from the AlphaFlow derived Alt1 and Alt2 models are β strand-like, and distinct from the ground state, for which ψ ∼ 0° is obtained (**Fig. 2C**). If Alt1 and Alt2 are models for ES_1_ and ES_2_, one would expect that replacing Q54 with a residue that favors β strand formation would stabilize the excited states. We prepared two such mutants of Q54, Q54V and Q54I (39), both of which increase the magnitude of dispersions observed in ^15^N CPMG experiments, suggesting that both variants have increased excited state populations (**fig. S3**). ^15^N CEST data were additionally recorded on Q54V (**table S2, S3**), the mutant with the largest CPMG dispersion profiles, and the combined analysis of ^15^N CPMG and CEST data yielded fractional populations of ES_1_ and ES_2_ of 6.4±0.2% and 7.6±0.3%, respectively, an increase of roughly 30-fold for ES_1_ and 20-fold for ES_2_ relative to the WT protein under the same experimental conditions (**Fig. 3A, table S4**). The larger populations and longer lifetimes of the Q54V variant excited states (8 ms and 16 ms for ES_1_ and ES_2_, respectively), increase the utility of CEST experiments, and, in contrast to the WT protein where minor state dips were only observed for ES_2_, the ^1^HN, ^15^N, ^13^CO, ^13^C^α^, ^13^C^β^, and ^13^C^methyl^ CEST profiles of some of the backbone and sidechain (^13^C^β^, ^13^C^methyl^) probes now show dips for both ES_1_ and ES_2_ in the Q54V mutant (**Fig. 3A**). Notably, the combined analysis of the Q54V ^15^N CPMG and CEST data required the use of a 4-state kinetic model (ES_3_↔GS↔ES_1_↔ES_2_) (**fig. S4**), indicating that this mutation increases the population of at least one other excited state as well. In addition, the chemical shift differences between the ground and ES_2_ states, Δν_ES2_ for the WT (x-axis) and Q54V (y-axis) proteins are similar (*r* = 0.89), suggesting that the Q54V mutation does not significantly perturb the structure of this excited state (**Fig. 3B**). Note that a full set of backbone chemical shifts was only available for ES_2_ of the WT protein (see supplementary materials; but available also for ES_1_ in the Q54V variant, **table S5**) so that a similar comparison involving ES_1_ is not possible. The correlation of Δν_ES2_ values is scattered, as expected, as the site of mutation is in the region where large chemical shift differences are found (**Fig. 1**, blue).

**Figure 3:**
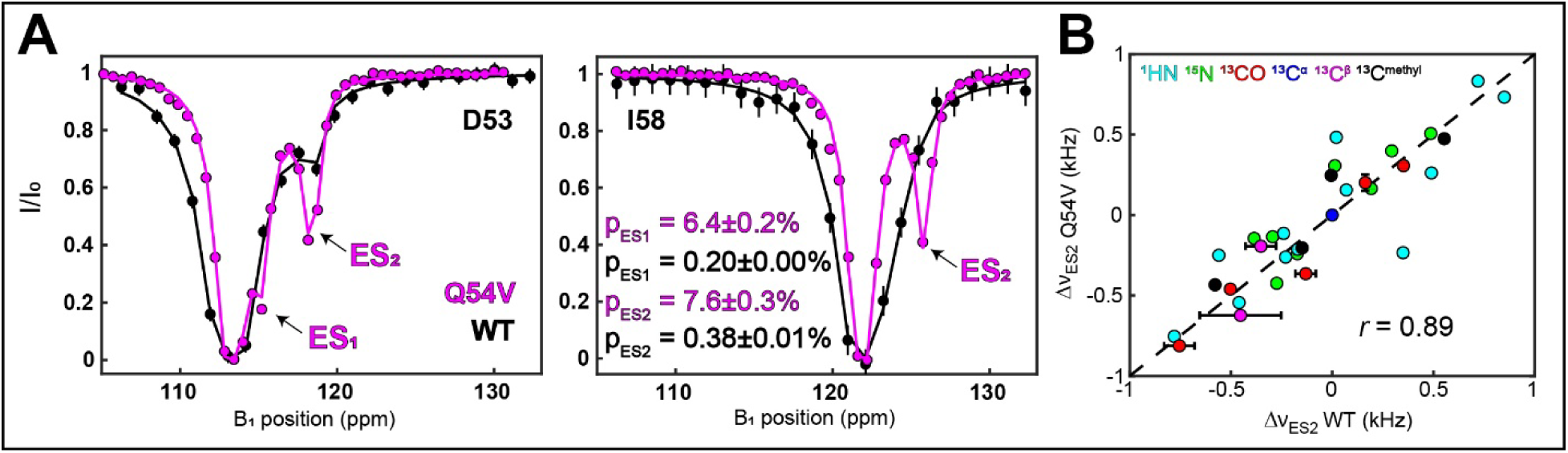
Replacing Q54 with a strand-favoring residue increases excited state populations. (A) ^15^N CEST profiles for D53 and I58. Profiles from the Q54V mutant (20 Hz spin lock, magenta) are overlaid with those from the WT (40 Hz spin lock, black), showing significantly more pronounced excited state dips for the mutant despite the lower spin lock power. The CEST profiles have been shifted so that the ground state dips overlap, and each point has been divided by exp(-*R_1_*T_CEST_) such that each baseline is at *I*/*I_0_* = 1 for ease of comparison. The Q54V data (4-state, ^15^N CPMG recorded at 1 GHz and 800 MHz and one ^15^N CEST dataset, see supplementary materials) are fit with populations for ES_1_ and ES_2_ that are more than an order of magnitude larger than those of the WT. (B) Chemical shift differences between ground and ES_2_ states of Q54V and WT (^1^HN, ^15^N, ^13^CO, ^13^C^α^, ^13^C^β^, ^13^C^methyl^) are in good agreement (*r* = 0.89), indicating that the mutation has not significantly changed the conformation of this excited state; corresponding shifts for ES_1_ are not available for the WT protein as the exchange kinetics preclude observation of CEST dips for this conformer (28). The Δν_ES2_ chemical shifts are given in kHz, calculated assuming a 1 GHz magnetic field. Chemical shifts of D53, Q/V54, and V55 are omitted from this comparison due to proximity to the site of mutation.

### Validation of AlphaFlow structure predictions by recording methyl-methyl NOEs

As the Q54V mutant increases the populations of ES_1_ and ES_2_ by over an order of magnitude relative to the WT protein, additional experiments can be recorded using it to validate the AlphaFlow-predicted conformers. The β* strand and its surroundings contain numerous Ile, Leu, and Val residues, and consequently there are several pairs of methyl groups which are only close together in Alt1 (red arrows), Alt2 (blue arrows), or in both (black arrows) where the β* strand is flipped around. This is shown schematically in **Figure 4A**, top and in the distograms below, highlighting the expected distances for some of the interactions based on analysis of the AlphaFlow predicted GS, Alt1, and Alt2 ensembles. Thus, observing NOEs between methyl groups that are predicted to be proximal in Alt1/Alt2 but not in the ground state in the Q54V mutant would provide evidence that Alt1 and Alt2 are structural models of the pair of excited states observed experimentally. This would then be further corroborated by the absence of such NOEs, or their significant reduction, in spectra recorded of the WT protein where the populations of ES_1_/ES_2_ are much smaller. **Figure 4B** shows a schematic of the 3D CCH NOESY experiment that was used in these studies, focusing on the magnetization transfer pathway that is most relevant for Alt1/Alt2 validation. During all three chemical shift encoding periods the signal of interest derives from the ground state. However, during the mixing time conversion from the ground to the excited states and subsequent build-up of NOEs within these lowly populated conformers occurs, as has been reported previously by Vallurupalli and coworkers (40). The cross peaks of interest, thus, report on interactions in the excited state, but are observed through ground state chemical shifts, providing a facile route to validate the alternative structures predicted by AlphaFlow.

**Figure 4:**
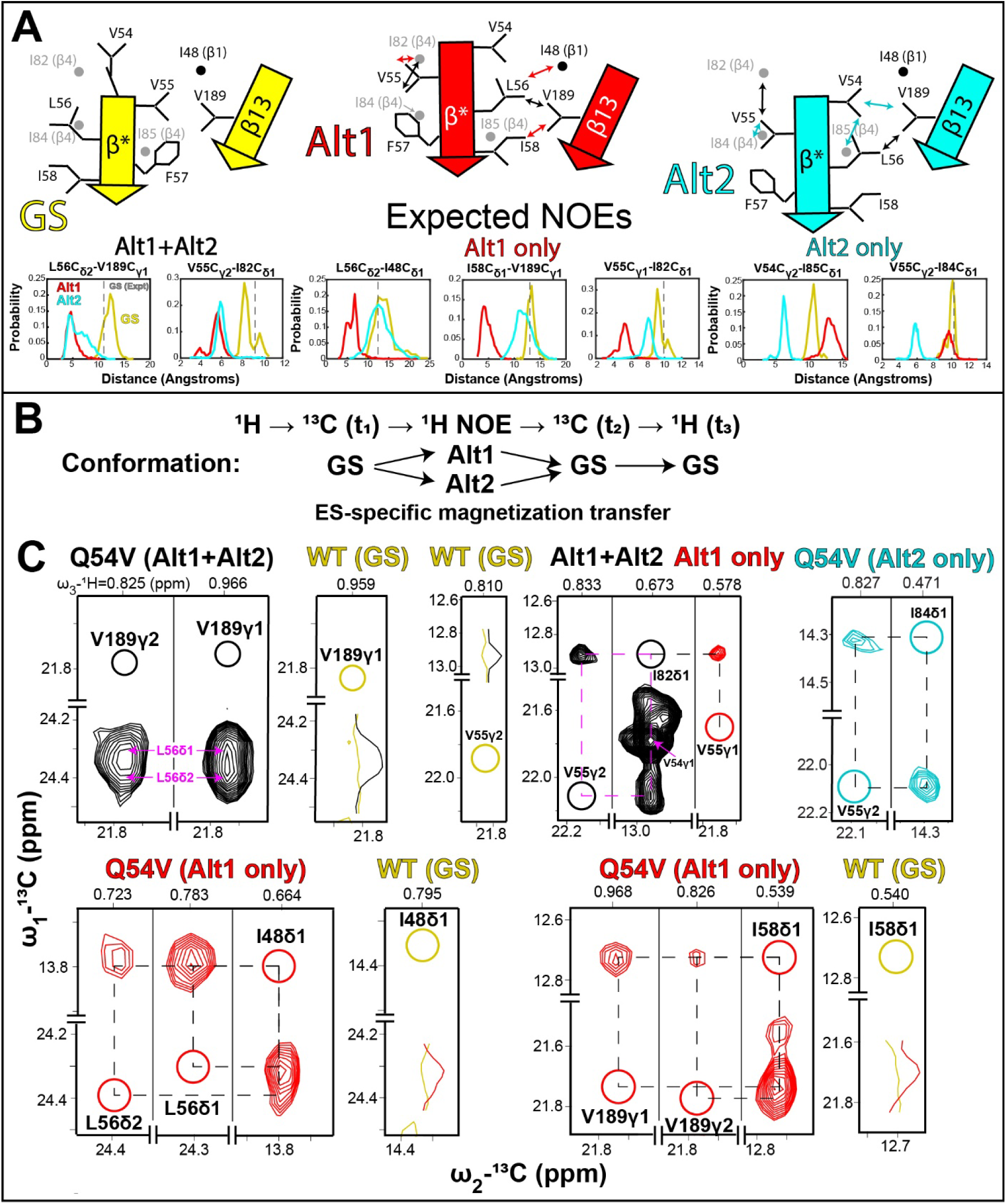
Validation of AlphaFlow structures via methyl-methyl NOEs. (A) (Top) Orientation of β* is shown for GS, Alt1, and Alt2 structures, with proximal methyl groups in either Alt1 (red), Alt2 (cyan), or both (black), but distant in GS, marked with arrows. Sidechains for residues from β* and β13 are indicated explicitly, while those from β1 and β4 are denoted by circles. (Bottom) Distance distributions between selected methyl groups amongst all Q54V AlphaFlow structures from the GS, Alt1, and Alt2 classes are highlighted. Dashed line indicates distance from GS, WT NMR structure (PDB 8URV) (46). (B) Schematic of the methyl-methyl CCH NOESY experiment, focusing on magnetization transfer steps that give rise to NOEs connecting proximal methyl groups within the excited states, observed through the ground state. (C) CCH NOESY slices for Q54V and WT pro-IL-18. NOEs derived from methyl groups close together in both Alt1 and Alt2 (but distant in GS), including L56δ1,2→V189γ1,2 (reciprocal cross peaks are observed but are overlapped with other signals) and V55γ2-I82δ1, give rise to cross peaks in the Q54V dataset (black) but not in the WT spectrum, or are much weaker (gold; compare black vs gold 1D traces in the adjacent WT slice). NOEs between proximal methyl groups only in Alt1, including I82δ1 →V55γ1, I48δ1-L56δ1,2 and I58δ1-V189γ1,2, lead to cross peaks in spectra recorded on samples of Q54V (red) but not of the WT (gold; compare red vs gold 1D traces in the proximal WT slice). 1D traces from the Q54V data set are shifted to align with the corresponding traces from the spectrum recorded on the WT protein. Also shown is a pair of NOESY peaks for Alt2 connecting V55γ2 with I84δ1 (plotted at a 30% lower level than all other slices). Diagonal peaks are not shown because of dynamic range issues but are indicated with circles and labeled to highlight the destination of magnetization transfer in the experiment.

CCH NOESY datasets were recorded on a 1 mM, highly deuterated, ILV-methyl labeled sample of Q54V pro-IL-18 (see Materials & Methods), as well as on an equivalently labeled and concentration matched WT sample as a control. There are five pairs of methyl groups which are expected to be close together in both Alt1 and Alt2 (Alt1+Alt2), but distant in the ground state, including both δ methyls of L56 and both γ methyls of V189, giving rise to four combinations of NOEs, and V55γ2-I82δ1 (**Fig. 4A**). In the CCH NOESY experiment recorded on the Q54V variant, cross peaks (weak relative to those originating from the ground state) are observed between all five of these methyl pairs, and three of these five were sufficiently resolved to be assigned (**Fig. 4C**). Importantly, there is comparatively very little to no signal observed between these methyl pairs in the WT dataset (**Fig. 4C**, yellow trace; **fig. S5**). This indicates that the NOESY cross peaks in the spectrum of Q54V originate from magnetization transfer in ES_1_ and/or ES_2_ and are not simply the result of spin diffusion within the ground state. Thus, L56δ1/2-V189γ1/2 and V55γ2-I82δ1 are close together in at least one of the excited states, validating the flip of the β* strand predicted by AlphaFlow. Notably, very weak cross peaks between L56δ1/2 and V189γ1 are observed in spectra recorded on the WT protein as well (**fig. S6**), further indicating that the Q54V excited states are structurally similar to those of the WT.

In addition, there are five pairs of methyl groups which are expected to be close only in Alt1, and distant in the ground state and Alt2. These include both δ methyls of L56 that are predicted to be proximal to I48δ1, I58δ1 that is proximal to both γ methyls of V189, and a V55γ1-I82δ1 contact (**Fig. 4A**, red arrows). In the CCH NOESY experiment recorded on the Q54V sample, cross peaks are observed between all five of these methyl pairs (**Fig. 4C**, red), with the corresponding cross peaks absent in the WT dataset (**Fig. 4C**, yellow trace; **fig. S5**). The presence of these cross peaks, along with those connecting L56δ1/2-V189γ1/2 and V55γ2-I82δ1, indicate that the Alt1 structure closely resembles one of the two experimentally observed excited states.

Several methyl pairs are also predicted to be proximal only in Alt2 and distant in both the ground state and Alt1, **Fig. 4A**. Although spectral overlap prevents a definitive assignment for many of these to distinct interactions, NOE contacts between V54γ2-I85δ1 (**fig. S5**) and V55γ2-I84δ1 (**Fig. 4C**) are observed that are telltale reporters of Alt2.

In addition to the NOE connectivities, other experimental data confirm important features of the Alt1/2 structures. For example, a TALOS analysis of ^1^HN, ^15^N, ^13^CO, ^13^C^α^, and ^13^C^β^ CEST-derived chemical shifts of ES_1_ and ES_2_ of the Q54V variant establish that residues V54-I58 are in β-sheet conformations, with the ψ of V54 transitioning from ∼ 0° in the ground state to ∼ 130° in both ES_1_ and ES_2_ (**table S1**). This change in ψ is a critical feature in the flip of the β* strand, providing strong evidence that both excited state conformations have similar flipped orientations and further validating the AlphaFlow structures. Additional evidence in support of the robustness of the Alt1 and Alt2 structures derives from a qualitative evaluation of *relative* intensities of NOEs. For example, NOEs connecting L56δ1/2 with V189γ1/2 are much stronger than the corresponding I58δ1-Val189γ1/2 correlations, as well as other Alt1/2 specific NOEs (**Fig. 4C, fig. S7**). This can be explained, at least qualitatively, by the AlphaFlow structures where residues 56 and 189 are close in both Alt1 and Alt2, while I58 and V189 are in similarly close proximity in Alt1 but distal in Alt2. Thus, both Alt conformers contribute to the L56-V189 NOE intensity, while only Alt1 is relevant for the I58-V189 NOE.

## DISCUSSION

The development of computational approaches for predicting protein structure has transformed the field of structural biology and it is now possible to generate three-dimensional models of biomolecules and their complexes with a high degree of confidence. However, computational approaches are not at the stage where they can predict biomolecular dynamics and energy landscapes of biomolecules with the same level of accuracy as ground state structures. Here we present a protocol which combines strengths of both NMR spin relaxation experiments and AlphaFlow to obtain validated structural models for sparsely populated and transiently formed protein conformers that are invisible to traditional biophysical experiments.

A combined CPMG and CEST NMR study establishes that pro-IL-18 interconverts between a ground state and a pair of excited states, enables quantification of the kinetics and thermodynamics of the exchange process, localizes the region of interconversion, and informs on the secondary structure of the exchanging regions in the sparse states. However, tertiary information is not forthcoming. AlphaFlow is complementary to the NMR approach because it produces a distribution of structures that can both be analyzed using existing NMR data and used for developing additional experiments to generate models of the excited states. In this manner a pair of AlphaFlow structures are identified, Alt1 and Alt2, as potential candidates. Both Alt structures show that Q54 becomes part of a β-sheet, distinct from the ground state and consistent with the NMR results, suggesting a Q54V variant to stabilize the excited states experimentally, as valine favors formation of β-strands more than glutamine (39). NMR studies of the Q54V protein indicate an increase in ES_1_ and ES_2_ populations by over an order of magnitude, so that NOE datasets can be recorded that validate both Alt1 and Alt2 as structural models of the pro-IL-18 excited states. The importance of NMR relaxation data in this process is underscored by the fact that the region undergoing exchange from ground to excited states needs to be masked in the generation of AlphaFlow conformers, since when provided the complete (*i.e.*, unmasked) structure as a template AlphaFlow was unable to produce structures other than the GS (**fig. S8**), and if a structural template is not provided correct structures of pro-IL-18 cannot be generated at all. The NMR data further provide an avenue to experimentally validate the utility of AlphaFlow, and more generally any generative computational model, for predicting invisible higher energy states on the energy landscape.

AlphaFlow generated models of the excited states of pro-IL-18 indicate that the β* strand is flipped relative to the ground state and that the primary difference in the excited state structures lies in a register shift of two amino acids in β*. β-strand flipping and/or slipping has been observed in a variety of proteins, including in ubiquitin (27,41), in mutants of the β-sheet OspA protein (42), and in protein-peptide complexes that are linked via inter-molecular β-sheets (43). In the case of pro-IL-18 these less stable configurations may play an important role in promoting unfolding of β* to prime binding of pro-IL-18 to caspases-1 and −4 for cleavage to the mature state of the protein.

Although the role of machine learning in structural biology will continue to advance rapidly (44), applications to studies of excited conformers will remain challenging, at least in the near future, due to the sparse data that is presently available. A combined NMR/machine learning strategy for studies of biomolecular dynamics (45) provides a powerful approach for exploring energy landscapes of increasing complexity, in ways that are not possible with either method alone, and ultimately will lead to the development of robust computational protocols for going beyond the prediction of static structure to include an accurate description of molecular dynamics.

## ACKNOWLEDGEMENTS

This work was supported through grants from the Canadian Institutes of Health Research (FND-503573) and the Natural Sciences and Engineering Council of Canada (024–03872) to L.E.K. J.B. acknowledges salary funding in the form of a post-doctoral fellowship from the CIHR.

## DATA AVAILABILITY

All AlphaFlow structural ensembles, Q54V and WT pro-IL-18 methyl-methyl CCH NOESY datasets, CPMG and CEST profiles for all analyzed probes, and TALOS input files for GS, ES_1_, and ES_2_ (Q54V) as well as ES_2_ (WT) are available on Zenodo (<u>10.5281/zenodo.17726228</u>). WT pro-IL-18 GS chemical shifts are available on the BMRB (31122).

## REFERENCES

1. Karplus M, Kuriyan J. Molecular dynamics and protein function. Proceedings of the National Academy of Sciences. 2005 May 10;102(19):6679–85.

2. Kurplus M, McCammon JA. Dynamics of Proteins: Elements and Function. Annu Rev Biochem. 1983 Jun;52(1):263–300.

3. Xie T, Saleh T, Rossi P, Kalodimos CG. Conformational states dynamically populated by a kinase determine its function. Science (1979). 2020 Oct 9;370(6513).

4. Hansen AL, Xiang X, Yuan C, Bruschweiler-Li L, Brüschweiler R. Excited-state observation of active K-Ras reveals differential structural dynamics of wild-type versus oncogenic G12D and G12C mutants. Nat Struct Mol Biol. 2023 Oct 28;30(10):1446–55.

5. Boehr DD, Mcelheny D, Dyson HJ, Wright PE. The Dynamic Energy Landscape of Dihydrofolate Reductase Catalysis. Science (1979). 2006;313(5793):1638–42.

6. Henzler-Wildman KA, Lei M, Thai V, Kerns SJ, Karplus M, Kern D. A hierarchy of timescales in protein dynamics is linked to enzyme catalysis. Nature. 2007 Dec 18;450(7171):913–6.

7. Whittier SK, Hengge AC, Loria JP. Conformational Motions Regulate Phosphoryl Transfer in Related Protein Tyrosine Phosphatases. Science (1979). 2013 Aug 23;341(6148):899–903.

8. Fraser JS, Clarkson MW, Degnan SC, Erion R, Kern D, Alber T. Hidden alternative structures of proline isomerase essential for catalysis. Nature. 2009 Dec 3;462(7273):669–73.

9. Neudecker P, Robustelli P, Cavalli A, Walsh P, Lundström P, Zarrine-Afsar A, et al. Structure of an Intermediate State in Protein Folding and Aggregation. Science (1979). 2012 Apr 20;336(6079):362–6.

10. Korzhnev DM, Religa TL, Banachewicz W, Fersht AR, Kay LE. A Transient and Low-Populated Protein-Folding Intermediate at Atomic Resolution. Science (1979). 2010 Sep 10;329(5997):1312–6.

11. Ceccon A, Tugarinov V, Ghirlando R, Clore GM. Abrogation of prenucleation, transient oligomerization of the Huntingtin exon 1 protein by human profilin I. Proceedings of the National Academy of Sciences. 2020 Mar 17;117(11):5844–52.

12. Baek M, DiMaio F, Anishchenko I, Dauparas J, Ovchinnikov S, Lee GR, et al. Accurate prediction of protein structures and interactions using a three-track neural network. Science (1979). 2021 Aug 20;373(6557):871–6.

13. Jumper J, Evans R, Pritzel A, Green T, Figurnov M, Ronneberger O, et al. Highly accurate protein structure prediction with AlphaFold. Nature. 2021 Aug 26;596(7873):583–9.

14. Ille AM, Anas E, Mathews MB, Burley SK. From sequence to protein structure and conformational dynamics with artificial intelligence/machine learning. Struct Dyn. 2025 May;12(3):030902.

15. Palmer AG, Kroenke CD, Patrick Loria J. Nuclear Magnetic Resonance Methods for Quantifying Microsecond-to-Millisecond Motions in Biological Macromolecules. In 2001. p. 204–38.

16. Carr HY, Purcell EM. Effects of Diffusion on Free Precession in Nuclear Magnetic Resonance Experiments. Physical Review. 1954 May 1;94(3):630–8.

17. Meiboom S, Gill D. Modified Spin-Echo Method for Measuring Nuclear Relaxation Times. Review of Scientific Instruments. 1958 Aug 1;29(8):688–91.

18. Vallurupalli P, Bouvignies G, Kay LE. Studying “invisible” excited protein states in slow exchange with a major state conformation. J Am Chem Soc. 2012 May 16;134(19):8148–61.

19. Bouvignies G, Vallurupalli P, Hansen DF, Correia BE, Lange O, Bah A, et al. Solution structure of a minor and transiently formed state of a T4 lysozyme mutant. Nature. 2011 Sep 21;477(7362):111–4.

20. Korzhnev DM, Vernon RM, Religa TL, Hansen AL, Baker D, Fersht AR, et al. Nonnative interactions in the FF domain folding pathway from an atomic resolution structure of a sparsely populated intermediate: an NMR relaxation dispersion study. J Am Chem Soc. 2011 Jul 20;133(28):10974–82.

21. Madhurima K, Nandi B, Munshi S, Naganathan AN, Sekhar A. Functional regulation of an intrinsically disordered protein via a conformationally excited state. Sci Adv. 2023 Jun 30;9(26).

22. Stiller JB, Otten R, Häussinger D, Rieder PS, Theobald DL, Kern D. Structure determination of high-energy states in a dynamic protein ensemble. Nature. 2022 Mar 17;603(7901):528–35.

23. Vallurupalli P, Hansen DF, Kay LE. Structures of invisible, excited protein states by relaxation dispersion NMR spectroscopy. Proceedings of the National Academy of Sciences. 2008 Aug 19;105(33):11766–71.

24. Fawzi NL, Ying J, Torchia DA, Clore GM. Probing exchange kinetics and atomic resolution dynamics in high-molecular-weight complexes using dark-state exchange saturation transfer NMR spectroscopy. Nat Protoc. 2012 Aug 19;7(8):1523–33.

25. Charlier C, Courtney JM, Anfinrud P, Bax A. Interrupted Pressure-Jump NMR Experiments Reveal Resonances of On-Pathway Protein Folding Intermediate. J Phys Chem B. 2018 Dec 13;122(49):11792–9.

26. Kitahara R, Yokoyama S, Akasaka K. NMR Snapshots of a Fluctuating Protein Structure: Ubiquitin at 30 bar–3 kbar. J Mol Biol. 2005 Mar;347(2):277–85.

27. Masoumzadeh E, Courtney JM, Charlier C, Ying J, Anfinrud P, Bax A. Structure of a transient protein-folding intermediate by pressure-jump NMR spectroscopy. Proceedings of the National Academy of Sciences. 2025 Oct 14;122(41).

28. Bonin JP, Aramini JM, Kay LE. Structural Plasticity as a Driver of the Maturation of Pro-Interleukin-18. J Am Chem Soc. 2024 Nov 6;146(44):30281–93.

29. Devant P, Dong Y, Mintseris J, Ma W, Gygi SP, Wu H, et al. Structural insights into cytokine cleavage by inflammatory caspase-4. Nature. 2023 Dec 14;624(7991):451–9.

30. Kaplanski G. Interleukin 18: Biological properties and role in disease pathogenesis. Immunol Rev. 2018 Jan 16;281(1):138–53.

31. Jing B, Berger B, Jaakkola T. AlphaFold Meets Flow Matching for Generating Protein Ensembles. ArXiv. 2024 Sep 2;

32. Lewis S, Hempel T, Jiménez-Luna J, Gastegger M, Xie Y, Foong AYK, et al. Scalable emulation of protein equilibrium ensembles with generative deep learning. Science (1979). 2025 Aug 14;389(6761).

33. Kalakoti Y, Wallner B. AFsample2 predicts multiple conformations and ensembles with AlphaFold2. Commun Biol. 2025 Mar 5;8(1):373.

34. Wayment-Steele HK, Ojoawo A, Otten R, Apitz JM, Pitsawong W, Hömberger M, et al. Predicting multiple conformations via sequence clustering and AlphaFold2. Nature. 2024 Jan 25;625(7996):832–9.

35. Lee M, Schafer JW, Prabakaran J, Chakravarty D, Clore MF, Porter LL. Large-scale predictions of alternative protein conformations by AlphaFold2-based sequence association. Nat Commun. 2025 Jul 1;16(1):5622.

36. Huang YJ, Ramelot TA, Spaman LE, Kobayashi N, Montelione GT. Hidden Structural States of Proteins Revealed by Conformer Selection with AlphaFold-NMR. bioRxiv. 2024.

37. Bonin JP, Aramini JM, Dong Y, Wu H, Kay LE. AlphaFold2 as a replacement for solution NMR structure determination of small proteins: Not so fast! Journal of Magnetic Resonance. 2024 Jul;364:107725.

38. Shen Y, Delaglio F, Cornilescu G, Bax A. TALOS+: a hybrid method for predicting protein backbone torsion angles from NMR chemical shifts. J Biomol NMR. 2009 Aug;44(4):213–23.

39. Regan L. Protein Structure: Born to be beta. Current Biology. 1994 Jul;4(7):656–8.

40. De D, Thapliyal N, Prakash Tiwari V, Toyama Y, Flemming Hansen D, Kay LE, et al. Mapping the FF domain folding pathway via structures of transiently populated folding intermediates. Proceedings of the National Academy of Sciences. 2024 Dec 10;121(50).

41. Gladkova C, Schubert AF, Wagstaff JL, Pruneda JN, Freund SM, Komander D. An invisible ubiquitin conformation is required for efficient phosphorylation by PINK1. EMBO J. 2017 Dec 15;36(24):3555–72.

42. Makabe K, Yan S, Tereshko V, Gawlak G, Koide S. β-Strand Flipping and Slipping Triggered by Turn Replacement Reveal the Opportunistic Nature of β-Strand Pairing. J Am Chem Soc. 2007 Nov 28;129(47):14661–9.

43. Panteva MT, Salari R, Bhattacharjee M, Chong LT. Direct Observations of Shifts in the β-Sheet Register of a Protein-Peptide Complex Using Explicit Solvent Simulations. Biophys J. 2011 May;100(9):L50–2.

44. Janson G, Feig M. Generation of protein dynamics by machine learning. Curr Opin Struct Biol. 2025 Aug;93:103115.

45. Wayment-Steele HK, El Nesr G, Hettiarachchi R, Kariyawasam H, Ovchinnikov S, Kern D. Learning millisecond protein dynamics from what is missing in NMR spectra. bioRxiv. 2025.

46. Dong Y, Bonin JP, Devant P, Liang Z, Sever AIM, Mintseris J, et al. Structural transitions enable interleukin-18 maturation and signaling. Immunity. 2024 Jul;57(7):1533–1548.e10.

