## Supplementary Materials for "Making invisible excited state protein structures visible by combining NMR and machine learning"

**Materials & Methods**

**Cloning, Expression, Purification, and Sample Preparation** **for NMR Studies**

Isotopically enriched samples of pro-IL-18 (WT, Q54I, and Q54V) were expressed and purified from *E. coli* using our previously established protocol (1). Briefly, the 193-amino acid coding sequence of pro-IL-18 was cloned into the Champion pET SUMO vector (ThermoFisher Scientific) in frame with an N-terminal SUMO and 6x His tag. The Q54 point mutations were introduced via site-directed mutagenesis using the Phusion high-fidelity PCR master mix (ThermoFisher Scientific), following the manual’s instructions for the PCR. Vectors were then transformed into *Escherichia coli* BL21 (DE3) cells and expressed in M9 minimal media containing either *U*-^15^NH_4_Cl and *U*-^13^C-glucose as the sole nitrogen and carbon sources for the production of *U*-[^13^C,^15^N] samples. For generating *U*-^2^H samples with ILV ^13^CH_3_-labeling, with both methyl groups of Leu and Val labeled (WT and Q54V; labeling of both isopropyl methyl groups is indicated by ^13^CH_3_/^13^CH_3_ in what follows), cells were grown in M9 D_2_O medium with *U*-^15^NH_4_Cl and d_7_-glucose, and precursors (60 mg/L α-ketobutyric acid, methyl-^13^C{3,3-D2} for Ileδ1-^13^CH_3_; 100 mg/L α-ketoisovaleric acid, dimethyl-^13^C2{3-D1} for Leuδ, Valγ-^13^CH_3_/^13^CH_3_) were added to media 1 hour prior to the induction of protein expression. The Leu/Val precursor was prepared by exchanging hydrogen with deuterium at the C3 position on the fully protonated molecule (α-ketoisovaleric acid, dimethyl-^13^C2). This was achieved via addition of 100 mg of the protonated precursor to 20 mL of 10 mM sodium phosphate at pD 12.5 in D_2_O, incubating at 45 ˚C and allowing exchange to proceed for 5 hours. Completion of the exchange reaction was confirmed by monitoring the decay of the corresponding ^1^H signal at the 3 position via 1D proton NMR.

Initial cell growths were carried out at 37 ^o^C, with induction of protein expression at 25 ^o^C by the addition of 0.1 mM IPTG at an OD_600_ of 0.8; expression continued for 20 hours. Proteins were purified at room temperature using 20 mM Tris buffers at pH 8 with 1 mM beta-mercaptoethanol. Cells were lysed by sonication on ice in the presence of 0.5 M KCl and a trace of DNase I. Clarified cell extracts were loaded onto a NiNTA column, purified by a wash step with 10 mM imidazole, and then the 6x His labeled protein was eluted using 0.3 M imidazole (0.1 M KCl). The 6x His and SUMO tags were then cleaved by Ulp protease during an overnight dialysis step to remove the imidazole, in the presence of 0.1 M KCl. A second NiNTA step was performed in which the tag-less pro-IL-18 was eluted in the flow-through. Final purification of pro-IL-18 was achieved via size exclusion chromatography using a prepacked 16/600 Superdex 75 column in the presence of 50 mM KCl. The final yield of purified isotopically-enriched pro-IL-18 was approximately 17 mg/L of culture for the WT, 11 mg/L of culture for the Q54V mutant, and 7 mg/L of culture for the Q54I mutant. Sample purity was confirmed by SDS-PAGE. Samples were concentrated by centrifugation to ~0.5-1 mM (see Methyl-Methyl NOESY section and Tables S2 and S3 below for concentrations of each sample) in 20 mM MES, 50 mM KCl, and 10 mM DTT at pH 6.5, 3% D_2_O, with 0.5 mM EDTA added to each sample.

**NMR Spectroscopy**

NMR experiments were collected at 25 ^o^C (unless stated otherwise) on Bruker AVANCE NEO 23.5 T (1.0 GHz), AVANCE III HD 18.8 T (800 MHz), and AVANCE III HD 14.1 T (600 MHz) NMR spectrometers equipped with 5-mm TCI triple-axis gradient cryoprobes. The recorded data were processed with NMRPipe(2), visualized using NMRFAM-SPARKY (3), and peak volumes were fit using peakipy (https://github.com/j-brady/peakipy). Our previously reported resonance assignments for pro-IL-18 were used (BMRB ID 31122) for the WT protein, and several additional experiments were recorded to facilitate the transfer of assignments to the Q54V spectrum (see Data Acquisition section below).

*Data Acquisition*

*Resonance Assignments of Q54V*

The assignments of many of the amide and methyl resonances of Q54V, as well as the amide resonances of Q54I, transferred readily from the WT spectrum. To assign amide resonances of Q54V which were ambiguous due to spectral crowding and/or large chemical shift perturbations with respect to the WT, we recorded HNCACB (4) and (HB/HA)CB/CA(CO)NH (5) datasets at 800 MHz on a 1 mM *U*-[^13^C,^15^N] sample. To assign ambiguous ILV methyl resonances, as well as to verify the assignments of all ILV methyl resonances in the Q54V variant, we recorded an H(CO)NH TOCSY dataset at 800 MHz on a 1 mM *U*-[^13^C,^15^N] sample, and an HMBC-HMQC (6) dataset at 800 MHz on a 1.2 mM *U*-[^2^H,^15^N] ILV ^13^CH_3_/^13^CH_3_ sample. Stereospecific assignments were generated by recording a CT-HSQC dataset (7,8) at 1 GHz on a 1 mM *U*-[5%-^13^C,^15^N] sample (9). Note that the Q54I variant was used only in initial CPMG and CEST experiments to compare relaxation profiles with those from the Q54V protein.

*CPMG and CEST*

^15^N, ^1^HN, and methyl CPMG (10–13) and CEST (14–16) experiments were recorded on *U*-[^2^H,^15^N] ILV ^13^CH_3_/^13^CH_3_ samples, while ^13^CO CPMG (17) and CEST (18), ^13^C^α^ CEST (19) and ^13^C^β^ CEST (20) experiments were recorded on *U*-[^13^C,^15^N] samples (see Tables S2 and S3 for further details). Spin lock carrier frequency sampling schedules for all CEST experiments were determined using an optimized frequency sampling approach (21). The ^1^HN CEST experiment was of the class whereby longitudinal order (IzSz, where Iz and Sz are z-components of ^1^H and ^15^N magnetization, respectively) is selected at the end of a relaxation period during which a weak B_1_ field is applied (22). Additional ^15^N CPMG experiments were recorded on *U*-[^15^N] samples of Q54V (25 ˚C, 40 ˚C), Q54I (40 ˚C), and WT (40 ˚C) pro-IL-18. For CPMG experiments, two duplicate planes were collected for peak intensity error estimation. Relevant parameters regarding the acquisition of the CPMG and CEST datasets are given in Tables S2 and S3, respectively.

*Methyl-methyl NOESY*

Methyl-methyl 3D CCH NOESY experiments were recorded on a 1.2 mM *U*-[^2^H,^15^N] ILV ^13^CH_3_ /^13^CH_3_ sample of the Q54V mutant and on a 1.1 mM *U*-[^2^H,^15^N] ILV ^13^CH_3_ /^13^CH_3_ sample of WT pro-IL-18. Both experiments were recorded with identical acquisition parameters, 250 ms mixing times, and 50% NUS (23). In addition, a 1 ms IBURP pulse (24) centered at 8 ppm was applied in the middle of the mixing element to invert the amide protons and thus minimize magnetization leakage to them during the mixing time (25).

*Data Analysis*

*CPMG and CEST*

All CPMG and CEST experiments were analyzed with the ChemEx program (https://github.com/gbouvignies/ChemEx). In fits of all CPMG and CEST data, transverse relaxation rates ($R_{2}$) of nuclei in the ground and excited states were constrained to be the same, unless a distinct excited state $R_{2}$ was required. For the backbone probes (^15^N, ^1^HN, ^13^CO), CPMG and CEST data for a given nucleus were analyzed together, constraining $\Delta\varpi$ and $R_{2}$ (at the same magnetic field) to be the same. ^15^N CPMG and CEST data from selected residues in β* and the preceding turn, including residues R49-V54 and F57-D59, were fit to a linear 4-state model (ES_3_↔GS↔ES_1_↔ES_2_) for the Q54V variant (details of fits of the relaxation data recorded on the WT protein were given previously (26)). The resulting populations and exchange rates were then fixed in the analysis of the ^1^HN data. For ^13^CO, ^13^C^α^, and ^13^C^β^ data, which were collected on protonated samples, slightly lower populations and exchange rates (see **table S4**) were used (relative to values optimal for deuterated samples), based on a linear 4-state fit of ^15^N CPMG and CEST data collected on a *U*-[^15^N] sample in which $\Delta\varpi_{AB}$, $\Delta\varpi_{AC}$, and $\Delta\varpi_{AD}$ were fixed to the values obtained from the analysis on the deuterated sample. For the sidechain methyl groups, the ^13^C CEST data were fit together with ^13^C MQ CPMG and ^1^H CEST profiles, with the populations and exchange rates fixed to the values obtained in the analysis of the backbone data.

**Generation of structural ensembles using AlphaFlow**

*Data Acquisition*

To generate the AlphaFlow (27) conformations of pro-IL 18 (both WT and Q54V) we used the 48-layer MD+template base version in which a template NMR structure of the protein is provided to the model, as we observed that the ground state structure of pro-IL18 was not predicted correctly without template inputs (28). We obtained multiple sequence alignments (MSAs) for pro-IL18 using ColabFold’s MSA server (29) and ran AlphaFlow in default settings. Notably, the full template did not generate any notable conformations that deviated from the input structure (**fig. S8**). Therefore, to allow diversification of the region of interest (L45-Q60), we masked the corresponding residues in the template by setting the atomic coordinates to zeros. A total of 30,000 conformations were generated across multiple NVIDIA P100s/V100s, corresponding to a computational cost of approximately 5 GPU days. A small number of structures were excluded from our analysis (50 for WT and 51 for Q54V) due to the presence of abnormally long bond distances.

*Data Analysis*

Backbone torsion angles of the 30,000 AlphaFlow structures were calculated (2 minutes) using IDPConformerGenerator’s torsions module (30) using 50 CPU threads. The criteria used to define the structural groups within the AlphaFlow datasets (**Fig. 2**) are as follows:

**GS**: Q/V54: -180˚ < φ < -75˚, -30˚ < ψ < 30˚; V55-I58: -180˚ < φ < -85˚, 100˚ < ψ < 180˚; Q/V54-I82 C^α^ – C^α^ distance < 5.5 Å; F57-I85 C^α^ – C^α^ distance < 5.5 Å

**Alt1**: Q/V54: -180˚ < φ < -75˚, 100˚ < ψ < 180˚; V55-I58: -180˚ < φ < -85˚, 100˚ < ψ < 180˚; Q/V54-T81 C^α^ –C^α^ distance < 7.5 Å; I58-I85 C^α^ –C^α^ distance < 5.5 Å

**Alt2**: Q/V54: -180˚ < φ < -75˚, 100˚ < ψ < 180˚; V55-I58: -180˚ < φ < -85˚, 100˚ < ψ < 180˚; Q/V54-F83 C^α^ –C^α^ distance < 5.5 Å; L56-I85 C^α^ –C^α^ distance < 5.5 Å

**Unfolded**: Q/V54: -180˚ < φ < -75˚, -30˚ < ψ < 30˚ or 100˚ < ψ < 180˚; V55-I58: -180˚ < φ < -85˚, 100˚ < ψ < 180˚ not all satisfied

**Other**: All other structures

A schematic illustrating the procedure used to classify the structures generated by AlphaFlow, highlighted in **Fig. 2B**, is shown in **fig. S2**. The criteria used to pick out structures were established based on an iterative approach, whereby NMR experiments informed analysis of the AlphaFlow results which, in turn, lead to further insights from the experimental data. For example, of the 30,000 structures produced for the WT protein, we initially focussed on those having φ,ψ dihedral values consistent with those for residues Q54-I58 of ES_2_ that were predicted by TALOS (31) using the ES_2_ chemical shifts obtained from the relaxation experiments (26). The low populations of ES_1_ and ES_2_ (< 0.5%), and the fact that no more than a single minor dip was observed in the CEST profiles recorded on the WT protein, precluded obtaining shifts for ES_1_, as described previously (26). The Alt1 structure was identified via manual inspection of a small number of the output structures at this stage and inspired the Q54V mutation that was predicted to increase the populations of the excited states. This was observed experimentally and notably two minor state dips were obtained in many of the CEST profiles of the Q54V variant so that φ,ψ dihedral values could be obtained also for residues V54-I58 in ES_1_, confirming that this segment is also a strand in this excited state. Subsequent analysis of the 3D CHH NOESY recorded on this variant indicated the possibility of an excited state involving a register shift by two amino acids relative to Alt1 (referred to as Alt2), as the relative intensities of L56-V189 NOEs (**fig. S7**) and Alt2 specific NOEs could not be explained by Alt1 alone (**Fig. 4C** and **fig. S5**). Based on the Alt1 structure and the predicted register shift defining a second conformation, as established from analysis of NOE experiments, C^α^ distances were chosen, as indicated above, to fix the register of the β* strand in either an Alt1 or Alt2 conformation. Structures were then grouped into ‘GS’, ‘Alt1’, ‘Alt2’, ‘Unfolded’, and ‘Other’ classes via the criteria highlighted above and, in this manner, the Alt2 structure, as predicted via the NOEs, was found in the ensemble of AlphaFlow-derived conformations. It is noteworthy that this change in register preserves hydrophobic contacts between β* and I48 (β1) and V189 (β13); hydrophobic contacts are also present in the GS and Alt1 conformations. In addition, these distance cutoffs ensure that the β* strand properly interacts with the β4 strand located below it, as we observed in some of the AlphaFlow structures that residues Q54-I58 were oriented roughly perpendicular to, rather than parallel to, the β4 strand (such structures belong to the ‘Other’ group). All these structural classes were also observed for the Q54V variant in AlphaFlow calculations.

**Supplementary Text**

**Backbone dihedral angles of β* in ES_1_ and ES_2_ from chemical shifts using TALOS**

In our previous study of WT pro-IL-18, we observed that CEST profiles contained no more than a single minor dip, corresponding to ES_2_, with minor dips from ES_1_ lacking (26). This can be explained by the kinetics of the exchange process, with the lifetime of the ES_1_ state sufficiently short to preclude its observation (26). Although CPMG data are informative in this situation, we were unable to record ^1^H^α^, ^13^C^α^, and ^13^C^β^ CPMG profiles with the *U*-[^13^C,^15^N] samples in hand due to interference from homonuclear scalar coupled evolution (32). Thus, only CEST datasets for ^1^H^α^, ^13^C^α^, and ^13^C^β^ probes were available that did not inform on ES_1_. In contrast, the exchange kinetics involving the GS, ES_1_, and ES_2_ states in the Q54V mutant are slowed and their populations significantly increased (over an order of magnitude, **Fig. 3**) so that distinct dips from both ES_1_ and ES_2_ are observed in cases where the chemical shifts of each of these states are distinct. In cases where the ES_1_, ES_2_, and GS ^13^C^α^ and ^13^C^β^ shifts are not unique, and only a single minor dip is observed in CEST profiles, it is not possible, in general, to determine whether the single dip derives from ES_1_ or ES_2_. Thus, backbone dihedral angles of the residues comprising β* were determined for both ES_1_ and ES_2_ using TALOS (**table S1**), considering only chemical shifts from Q54V pro-IL-18 which could be confidently determined for ES_1_ and ES_2_ (**table S5**). The chemical shift analysis reveals that the β* strand is flipped in both ES_1_ and ES_2_, based on the change in ψ of V54 (Q54 in WT). Notably, the backbone dihedral angle predictions for β* residues in ES_2_ for the Q54V variant are nearly identical to those for the corresponding region of ES_2_ in the WT protein (**table S1**), as expected given the similarity in the chemical shifts (**Fig. 3B**).

**A 4-state kinetic model is necessary to globally fit Q54V ^15^N CPMG and CEST data**

Initial attempts to fit the ^15^N CPMG and CEST data from the Q54V mutant with a linear 3-state model (GS↔ES_1_↔ES_2_), as we have previously used for the WT protein (26), showed that although the CEST data could be reproduced very well, the model could not simultaneously replicate the CPMG data (**fig. S4**). One possibility is an additional faster process (or processes) that contributes to the CPMG but not to the CEST profiles. Consequently, we fit the Q54V CPMG and CEST data to a linear four state model (ES_3_↔GS↔ES_1_↔ES_2_) to include a faster interconversion of GS with an additional state, ES_3_. This simple addition to the original 3-state model yields significantly improved fits to the CPMG data without compromising the quality of the fits of the CEST data, as is shown for D53 and V54 in **fig. S4**. Given the rapid exchange between GS and ES_3_, the population of ES_3_ cannot be determined. The best fit value of this population is 0.7%, but nearly identical fits are obtained when fixing the population of ES_3_ at much higher values.

**Supplementary Figures**

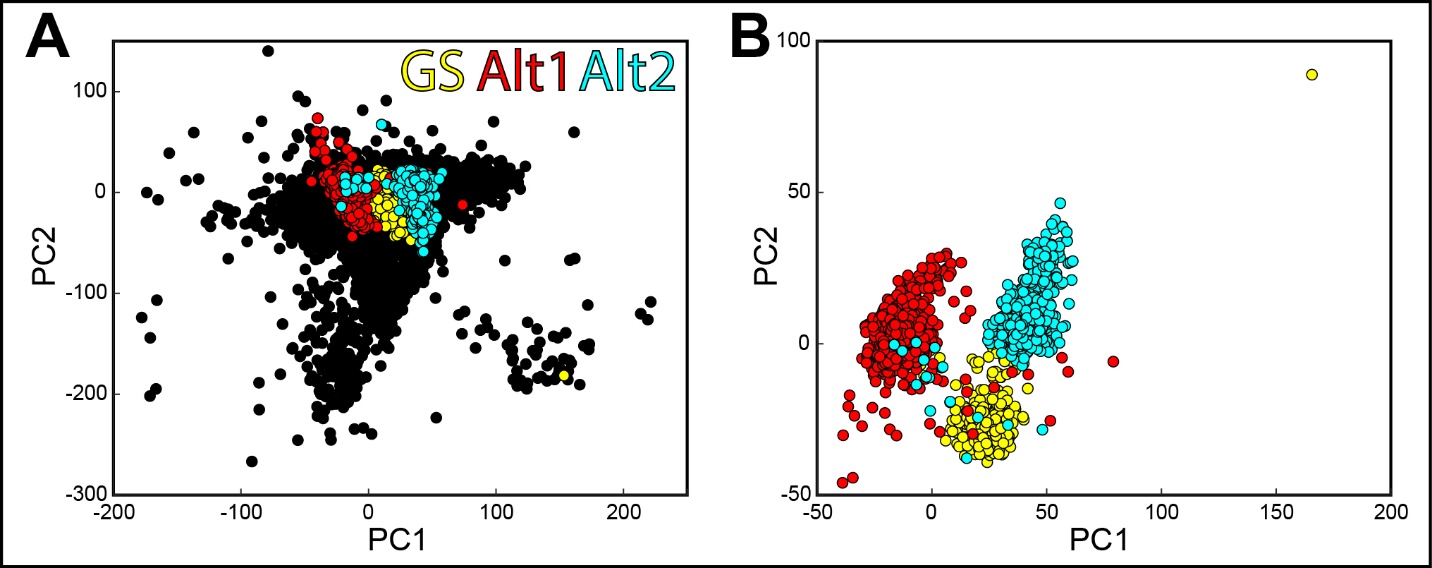

**Fig. S1: Principal component analysis is unable to identify GS, Alt1, and Alt2 groups when using all AlphaFlow structures.** (A) Principal component analysis of atomic coordinates (residues 45-60) from all AlphaFlow structures for WT pro-IL-18. Data points from structures in the GS, Alt1, and Alt2 groups are colored in yellow, red, and cyan, respectively. Data points from all other structures (Unfolded, Other) are colored in black. When including all structures, the GS, Alt1, and Alt2 groups are obscured by a single, large cluster of structures that likely results from the variability of atomic coordinates within the Unfolded group. (B) Principal component analysis of atomic coordinates (residues 45-60) from only the structures within the GS, Alt1, and Alt2 groups. After removing the structures from the Unfolded and Other groups, a clear separation of the GS, Alt1, and Alt2 structures is observed.

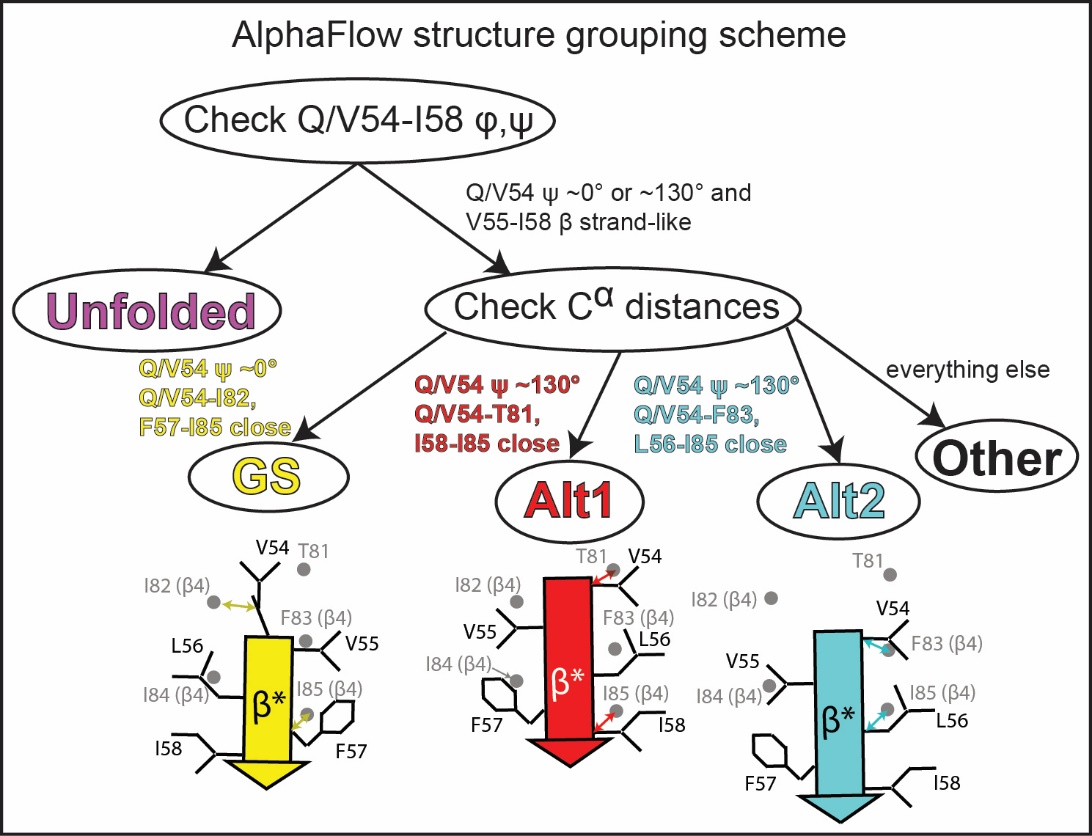

**Fig. S2: Procedure for assigning AlphaFlow conformers to structural groups.** Schematic showing how AlphaFlow conformers were separated into five structural groups, including GS, Alt1, and Alt2. The rationale for the various criteria used is described in the Materials & Methods. Also highlighted are cartoon representations of β* in GS, Alt1, and Alt2, with arrows to highlight the C^α^-C^α^ contacts that are used to distinguish the different conformers.

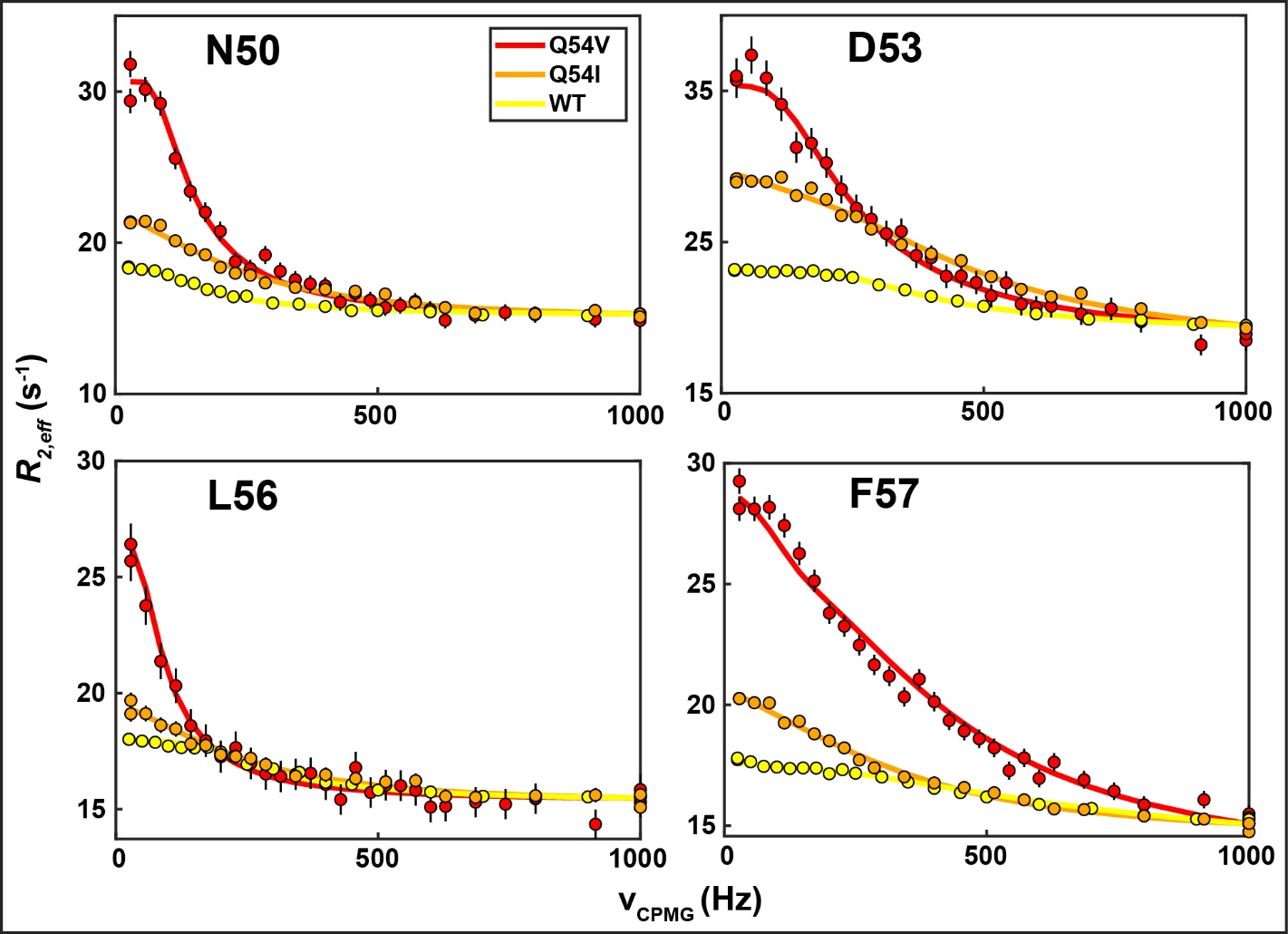

**Fig. S3: Mutation of Q54 to β strand-favoring residues increases excited state populations.** ^15^N CPMG profiles for selected residues are shown for Q54V (red), Q54I (orange), and WT (yellow) pro-IL-18. Datasets were collected under identical conditions (800 MHz, 40 ˚C, see **table S2**). Increases in dispersion magnitudes are observed for both Q54V and Q54I relative to WT, suggesting that replacing Q54 with a β strand-favoring residue such as valine or isoleucine provides increased stabilization of the excited states. The Q54V mutant was selected for further study, as the dispersions are considerably larger for this mutant compared to Q54I.

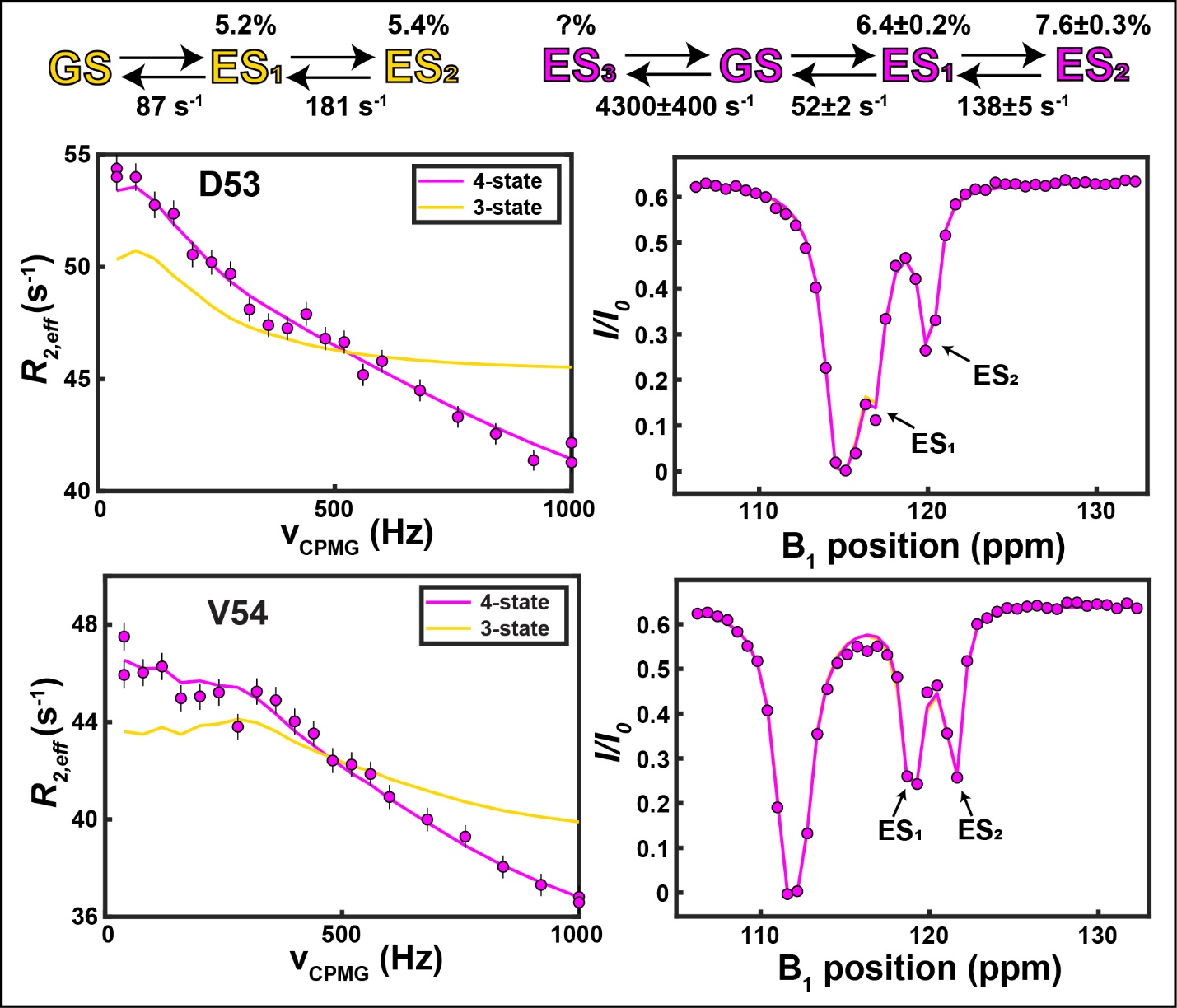

**Fig. S4: A 4-state kinetic model is required to fit Q54V ^15^N CPMG and CEST data.** While the linear 3-state model (GS↔ES_1_↔ES_2_) used to fit the CPMG and CEST data recorded on WT pro-IL18 in our previous work (26) can accurately reproduce the Q54V CEST data, it yields very poor fits of the corresponding CPMG data. A 4-state model having an additional excited state connected only to the ground state (ES_3_↔ GS↔ES_1_↔ES_2_) can fit both the CPMG and CEST profiles, with similar exchange parameters for ES_1_ and ES_2_ compared to the 3-state model. Given the rapid exchange between GS and ES_3_, the population of ES_3_ cannot be determined.

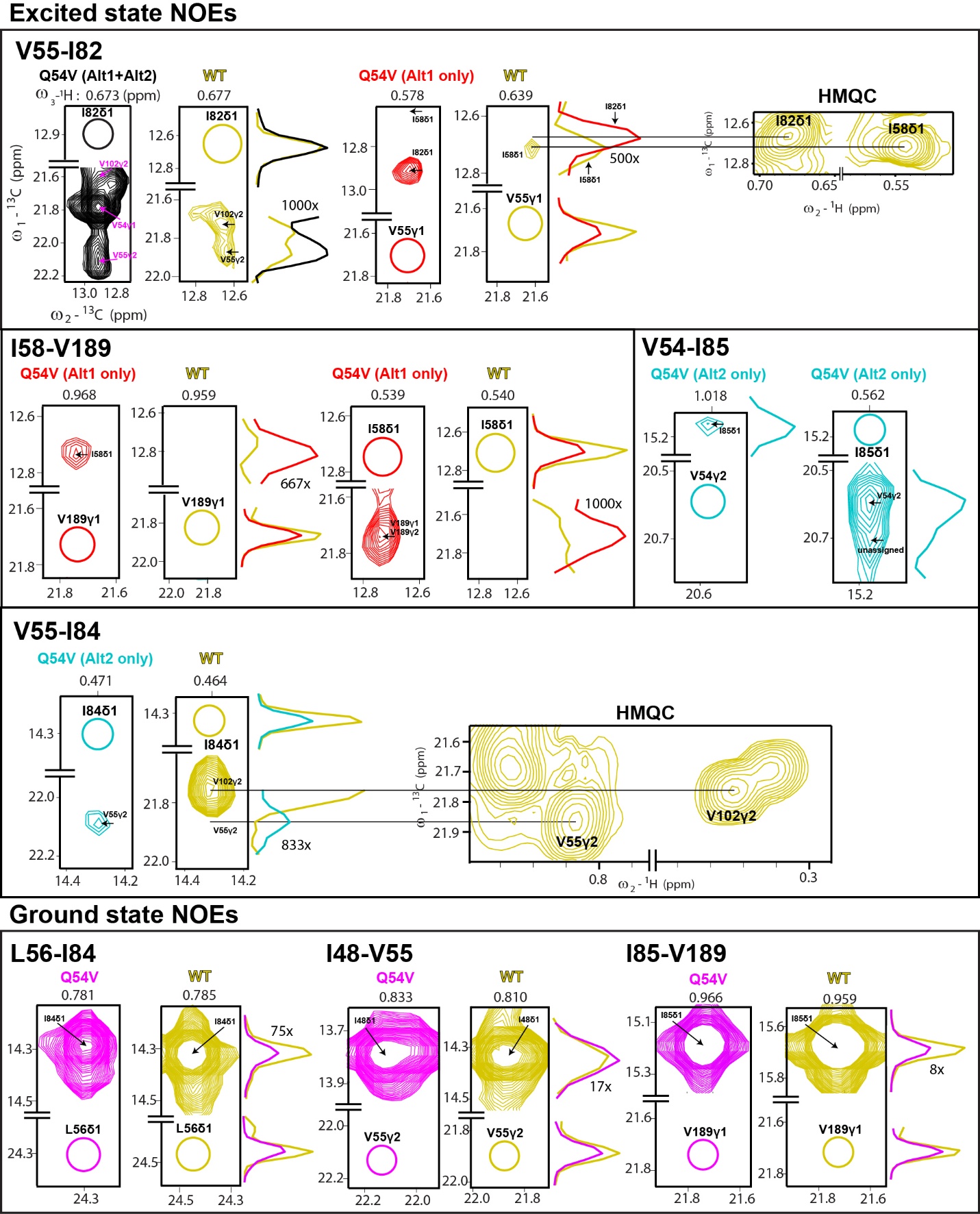

**Fig. S5: Q54V NOESY data contain excited state-specific cross peaks which are absent (or much weaker) in WT.** Comparisons of selected NOESY slices/traces from data recorded on Q54V and WT proteins are shown for all unambiguous excited state-specific NOEs that are not highlighted in **Fig. 4**, as well as for the V54-I85 NOEs which are not present in spectra of the WT due to the absence of Val at position 54, but which are expected to be specific to the Alt2 conformation based on distances predicted from the AlphaFlow-derived structures (**Fig. 4**, top). Traces from the Q54V dataset shown to the sides of the NOE panels have been shifted to align with either the corresponding peaks from the WT data or their expected positions (yellow) for ease of visualization (compare yellow and black traces corresponding to I82δ1 diagonal peaks for V55-I82 NOEs, top row, for example). Traces of cross peaks (not diagonal peaks) have been scaled for visualization, with the scaling factor indicated. All data are plotted at the same starting contour level (1.0e11), with the exception of the pair of slices showing the I82δ1-V55γ1 peaks (top row) that are displayed starting at a threshold that is 30% lower (0.7e11) so as to visualize the cross peak with I58δ1 in the data of the WT protein. Excited state NOEs (cross peaks) are generally not observed in data recorded on WT pro-IL-18, despite the fact that the diagonal peaks from the WT protein are generally more intense than the corresponding peaks in spectra of the Q54V variant. Similarly, NOEs originating from the ground state (bottom) are generally more intense in NOESY spectra of the WT or at least as intense as in corresponding spectra recorded on Q54V.

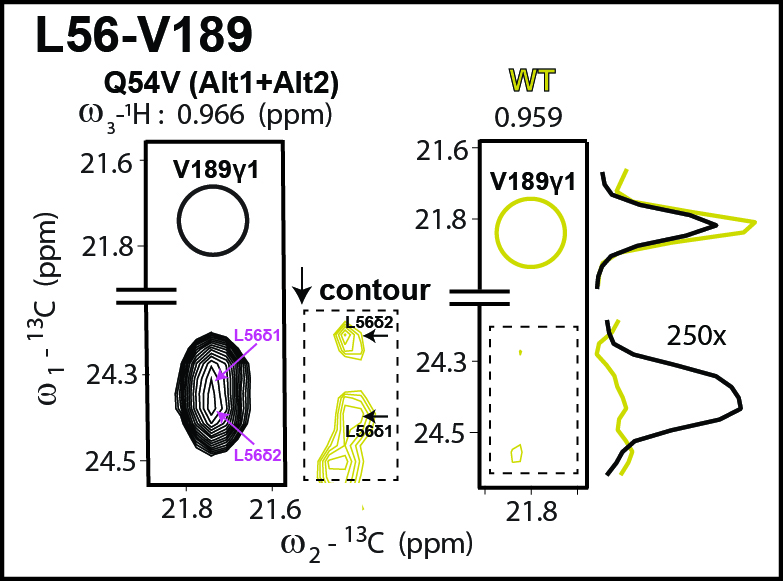

**Fig. S6: L56δ1,2-V189γ1 cross peaks are observed in the WT protein, albiet at low intensity.** Cross peaks between L56δ1,2 and V189γ1, consistent with both Alt1 and Alt2 conformations and indicative of the flip of the β* strand in the excited states, are observed in NOESY spectra of the WT protein, although they are much weaker than in spectra recorded of the Q54V mutant. The contour level of the inset is 0.7e11, while the other panels have a contour level of 1.0e11. This suggests that the excited states of the Q54V mutant are similar to those of the WT protein, consistent with the similarity in ES_2_ chemical shifts between Q54V and WT pro-IL-18 (**Fig. 3B**).

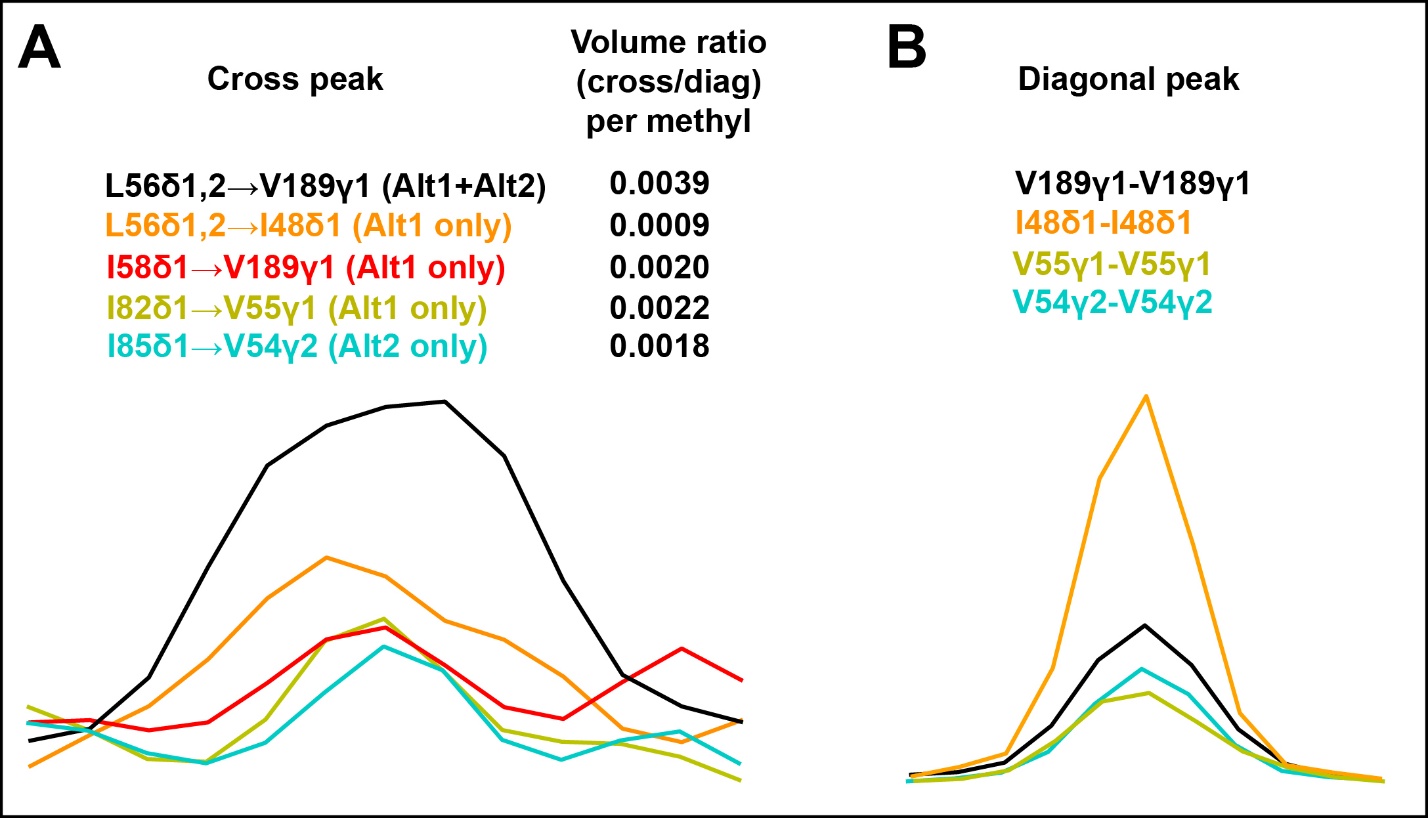

**Fig. S7: Relative volume of the L56δ1,2→V189γ1 cross peak is consistent with contributions from both Alt1 and Alt2.** Volume ratios are calculated by dividing the peak volume of the cross peak (A) by that of the corresponding diagonal peak (B). In the case of the L56δ1,2→V189γ1 and L56δ1,2→I48δ1 cross peaks, corresponding to magnetization transfer from L56 to V189 or I48 respectively (as indicated by the arrows), which have contributions from both methyl groups of L56, the volume ratio is divided by two (note that separate NOEs from L56δ1 and δ2 are not observed, as the ^13^Cδ shifts are nearly degenerate). The normalized volume ratio of the L56δ1,2→V189γ1 cross peak is roughly twice as large as that of the I58δ1→V189γ1, I82δ1→V55γ1, and I85δ1→V54γ2 cross peaks. The difference is larger, roughly four-fold, with respect to the L56δ1,2→I48δ1 cross peak. This is consistent with L56 and V189 being proximal in both ES_1_ and ES_2_, while the other pairs of methyl groups are proximal in only one of the two excited states. The Alt1 and Alt2 structures predicted by AlphaFlow satisfy these criteria, consistent with the notion that Alt1 and Alt2 resemble the two excited states.

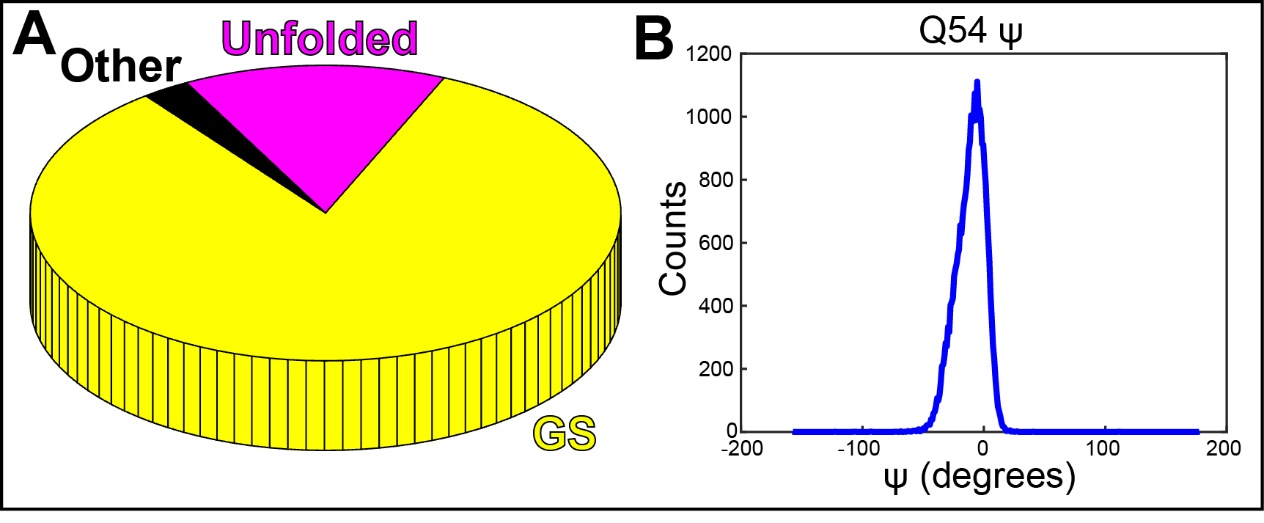

**Fig. S8: AlphaFlow does not generate the Alt1 and Alt2 structures without masking the β* region.** 30,000 structures of WT pro-IL-18 were generated by AlphaFlow using our experimental structure as a template with no masking. (A) Fraction of conformers from the five structural groups are illustrated as a pie chart. Without masking, the majority of the structures generated belong to the GS group (*i.e.* they are very similar to the template, experimental structure). Not a single Alt1 or Alt2 structure was generated. (B) A histogram of the Q54 ψ angle from this set of structures shows a single mode near 0˚ (GS), with no structures near 130˚ (Alt1/Alt2). This emphasizes the importance of masking the region of interest to obtain diverse conformations from the starting structure.

**Supplementary Tables**

| State | Q/V54 φ | Q/V54 ψ | V55 φ | V55 ψ | L56 φ | L56 ψ | F57 φ | F57 ψ | I58 φ | I58 ψ |
| --- | --- | --- | --- | --- | --- | --- | --- | --- | --- | --- |
| Q54V GS^a^ | -95±13 | -3±19 | -122±23 | 133±19 | -125±25 | 131±16 | -109±11 | 125±18 | -114±19 | 132±16 |
| Q54V GS^b^ | -95±16 | -12±14 | -127±19 | 132±11 | -121±26 | 131±12 | -114±15 | 128±20 | -124±20 | 144±14 |
| WT GS^c^ | -99±15 | 1±16 | -118±22 | 136±17 | -124±16 | 138±7 | -112±15 | 133±14 | -115±22 | 131±14 |
| Q54V ES_1_^d^ | -127±21 | 131±13 | -116±16 | 128±12 | -100±14 | 123±7 | -113±19 | 137±14 | -110±22 | 143±15 |
| Q54V ES_2_^e^ | -121±22 | 134±14 | -112±19 | 130±12 | -110±14 | 128±11 | -119±13 | 125±16 | -113±14 | 121±9 |
| WT ES_2_^c^ | -68±65 | 116±32 | -136±24 | 146±21 | -117±21 | 122±9 | -123±13 | 134±13 | -112±12 | 125±12 |

**Table S1: Backbone dihedral angles of β* in GS, ES_1_, and ES_2_ from TALOS** (31)

^a^ ^15^N, ^1^HN, ^13^CO, ^13^C^α^, ^13^C^β^, and ^1^H^α^ shifts used for D53-I58 and ^15^N, ^1^HN, ^13^C^α^, and ^13^C^β^ shifts used for D59 (^13^CO and ^1^H^α^ not assigned).

^b^ Including only shifts for probes (nuclei) that are also available for ES_1_ and ES_2_ (see **table S5**).

^c 15^N, ^1^HN, ^13^CO, ^13^C^α^, ^13^C^β^, and ^1^H^α^ shifts used for all residues.

^d^ Based on shifts available for ES_1_ (**table S5**).

^e^ Based on shifts available for ES_2_ (**table S5**).

Orange background indicates that the dihedral angle was predicted with low confidence.

| Nucleus: (Q54V/Q54I/WT) | Temp (^o^C) | [pro-IL-18] (mM) | B_0_ (MHz) | CPMG Relaxation time (ms) | $\nu_{CPMG}$ range (Hz) |
| --- | --- | --- | --- | --- | --- |
| ^15^N: Q54V | 25 | 1.2 | 1000, 800 | 25 | 40 – 1000 (20 values) |
| ^15^N: (*U*-[^15^N]) Q54V | 25 | 1.0 | 800 | 30 | 33 – 1000 (20 values) |
| ^15^N: (*U*-[^15^N]) Q54V | 40 | 0.8 | 800 | 35 | 29 – 1000 (27 values) |
| ^15^N: (*U*-[^15^N]) Q54I | 40 | 0.65 | 800 | 35 | 29 – 1000 (20 values) |
| ^15^N: (*U*-[^15^N]) WT | 40 | 1.1 | 800 | 40 | 25 – 1000 (20 values) |
| ^1^HN: Q54V | 25 | 1.2 | 1000 | 20 | 50 – 2000 (19 values) |
| ^13^CO: Q54V | 25 | 0.6 | 800 | 20 | 50 – 950 (19 values) |
| Methyl ^13^C: Q54V | 25 | 1.2 | 1000 | 25 | 40 – 2000 (20 values) |

**Table S2: Acquisition parameters of CPMG datasets.**

| Nucleus | Temp (^o^C) | [pro-IL-18] (mM) | B_0_ (MHz) | Relaxation time (ms) | Weak B_1_ field (Hz) | Frequency range (ppm) |
| --- | --- | --- | --- | --- | --- | --- |
| ^15^N | 25 | 1.2 | 800 | 500 | 20.9 | 106 – 132 (45 values) |
| ^15^N (*U*-[^15^N]) | 25 | 1.0 | 800 | 500 | 20.9 | 106 – 132 (45 values) |
| ^1^HN | 25 | 1.2 | 800 | 400 | 19.5 | 6.5 – 10.7 (70 values) |
| ^13^CO | 25 | 0.6 | 600 | 300 | 26.4 | 169 – 181 (30 values) |
| ^13^C^α^ | 25 | 1.2 | 800 | 500 | 27.2 | 54 – 65  (47 values) |
| ^13^C^β^ | 25 | 1.2 | 800 | 350 (Val, Ile), 300 (Leu) | 27.2 | 29.5 – 37 (32 values, Val)  37.3 – 43.3  (26 values, Ile)  41.5-49.5  (28 values, Leu) |
| Methyl ^13^C | 25 | 1.2 | 800 | 350 | 21.8 | 9.7 – 28.3 (78 values) |
| Methyl ^1^H | 25 | 1.2 | 800 | 300 | 19.5 | 0.0 – 1.4  (24 values) |

**Table S3: Acquisition parameters of CEST datasets on Q54V pro-IL-18.**

| Sample | $p_{ES1}(\%)$ | $p_{ES2}(\%)$ | $p_{ES3}$ | $k_{ex,GSES1}(s^{-1})$ | $k_{ex,ES1ES2}(s^{-1})$ | $k_{ex,GSES3}(s^{-1})$ |
| --- | --- | --- | --- | --- | --- | --- |
| *U*-[^2^H,^15^N] ILV ^13^CH_3_ /^13^CH_3_ | $6.4\pm0.2$ | $7.6\pm0.3$ | N/D | $52\pm2$ | $138\pm5$ | $4300\pm400$ |
| *U*-[^15^N] | $5.7\pm0.4$ | $5.8\pm0.5$ | N/D | $48\pm4$ | $138\pm10$ | $2600\pm200$ |

**Table S4: 4-state model exchange parameters for ^2^H and ^1^H Q54V samples.**

| Residue | ^1^HN $\Delta\varpi_{GS,ES1}$ | | ^1^HN $\Delta\varpi_{GS,ES2}$ |
| --- | --- | --- | --- |
| F13 | $-0.749\pm0.003$ | | $-0.114\pm0.001$ |
| L23 | $-0.545\pm0.003$ | | $-0.234\pm0.003$ |
| I48 | $-0.207\pm0.002$ | | $0.156\pm0.002$ |
| R49 | $0.2098\pm0.0008$ | | $-0.213\pm0.001$ |
| N50 | $-0.063\pm0.002$ | | $-0.2611\pm0.0007$ |
| L51 | $0.4477\pm0.0006$ | | $0.8327\pm0.0007$ |
| D53 | $0.045\pm0.006$ | | $-0.586\pm0.001$ |
| V54 | $-0.168\pm0.001$ | | $0.619\pm0.002$ |
| V55 | $1.4414\pm0.0008$ | | $0.4925\pm0.0007$ |
| L56 | $-0.426\pm0.001$ | | $-0.544\pm0.001$ |
| F57 | $0.4406\pm0.0005$ | | $0.7325\pm0.0008$ |
| I58 | $-0.9068\pm0.0008$ | | $-0.754\pm0.001$ |
| D59 | $-0.6061\pm0.0006$ | | $0.2615\pm0.0009$ |
| N62 | $0.2417\pm0.0006$ | | $0.4830\pm0.0007$ |
| Q190 | $0.066\pm0.003$ | | $-0.250\pm0.001$ |
| Residue | | ^15^N $\Delta\varpi_{GS,ES1}$ | ^15^N $\Delta\varpi_{GS,ES2}$ |
| R49 | | $-0.39\pm0.03$ | $-4.18\pm0.01$ |
| N50 | | $-1.71\pm0.02$ | $-2.37\pm0.01$ |
| L51 | | $-1.96\pm0.02$ | $3.02\pm0.01$ |
| D53 | | $1.86\pm0.02$ | $5.20\pm0.02$ |
| V54 | | $7.06\pm0.01$ | $9.66\pm0.02$ |
| V55 | | $8.44\pm0.02$ | $7.09\pm0.06$ |
| L56 | | $-1.13\pm0.06$ | $-1.43\pm0.02$ |
| F57 | | $2.131\pm0.009$ | $4.996\pm0.009$ |
| I58 | | $-0.46\pm0.04$ | $3.93\pm0.02$ |
| D59 | | $-1.77\pm0.02$ | $1.62\pm0.02$ |
| I82 | | $-0.37\pm0.03$ | $-1.33\pm0.01$ |

| Residue | ^13^CO $\Delta\varpi_{GS,ES1}$ | ^13^CO $\Delta\varpi_{GS,ES2}$ |
| --- | --- | --- |
| I48 | $-0.1\pm0.2$ | $-1.45\pm0.06$ |
| R49 | $0.0\pm0.2$ | $-1.83\pm0.05$ |
| D53 | $0.0\pm0.2$ | $-2.49\pm0.05$ |
| V54 | $1.1\pm0.1$ | $1.4\pm0.1$ |
| V55 | $-1.13\pm0.06$ | $-1.65\pm0.05$ |
| L56 | $2.05\pm0.03$ | $1.22\pm0.07$ |
| F57 | $-1.82\pm0.04$ | $-3.23\pm0.04$ |
| I58 | $0.92\pm0.08$ | $0.8\pm0.2$ |

| Residue | ^13^C^α^ $\Delta\varpi_{GS,ES1}$ | ^13^C^α^ $\Delta\varpi_{GS,ES2}$ |
| --- | --- | --- |
| F57 | $0$ | $0$ |

| Residue | ^13^C^β^ $\Delta\varpi_{GS,ES1}$ | ^13^C^β^ $\Delta\varpi_{GS,ES2}$ |
| --- | --- | --- |
| V55 | $0.87\pm0.02$ | $-1.15\pm0.02$ |
| L56 | $-2.688\pm0.008$ | $-0.77\pm0.02$ |
| I58 | $-0.80\pm0.02$ | $-2.48\pm0.01$ |

| Methyl Group | ^13^C $\Delta\varpi_{GS,ES1}$ | ^13^C $\Delta\varpi_{GS,ES2}$ |
| --- | --- | --- |
| L51δ2 | $-0.827\pm0.003$ | $0.980\pm0.003$ |
| V55γ2 | $-0.836\pm0.005$ | $-1.724\pm0.004$ |
| L56δ2 | $0.663\pm0.003$ | $1.886\pm0.003$ |
| I85δ1 | $-0.485\pm0.003$ | $-0.812\pm0.002$ |

**Table S5: Chemical shift differences (ppm) between ground and excited states (ES_1_ and ES_2_) for Q54V mutant (**$\boldsymbol{\Delta}\boldsymbol{\varpi}_{\boldsymbol{A,B}}\boldsymbol{=}\boldsymbol{\varpi}_{\boldsymbol{B}}\boldsymbol{-}\boldsymbol{\varpi}_{\boldsymbol{A}}\boldsymbol{)}$**.**

**References**

1.	Dong Y, Bonin JP, Devant P, Liang Z, Sever AIM, Mintseris J, et al. Structural transitions enable interleukin-18 maturation and signaling. Immunity. 2024 Jul;57(7):1533-1548.e10.

2.	Delaglio F, Grzesiek S, Vuister GeertenW, Zhu G, Pfeifer J, Bax A. NMRPipe: A multidimensional spectral processing system based on UNIX pipes. J Biomol NMR. 1995 Nov;6(3).

3.	Lee W, Tonelli M, Markley JL. NMRFAM-SPARKY: enhanced software for biomolecular NMR spectroscopy. Bioinformatics. 2015 Apr 15;31(8):1325–7.

4.	Wittekind M, Mueller L. HNCACB, a High-Sensitivity 3D NMR Experiment to Correlate Amide-Proton and Nitrogen Resonances with the Alpha- and Beta-Carbon Resonances in Proteins. J Magn Reson B. 1993 Apr;101(2):201–5.

5.	Grzesiek S, Bax A. Correlating backbone amide and side chain resonances in larger proteins by multiple relayed triple resonance NMR. J Am Chem Soc. 1992 Jul 1;114(16):6291–3.

6.	Siemons L, Mackenzie HW, Shukla VK, Hansen DF. Intra-residue methyl–methyl correlations for valine and leucine residues in large proteins from a 3D-HMBC-HMQC experiment. J Biomol NMR. 2019 Dec 12;73(12):749–57.

7.	Santoro J, King GC. A constant-time 2D overbodenhausen experiment for inverse correlation of isotopically enriched species. Journal of Magnetic Resonance (1969). 1992 Mar;97(1):202–7.

8.	Vuister GW, Bax A. Resolution enhancement and spectral editing of uniformly 13C-enriched proteins by homonuclear broadband 13C decoupling. Journal of Magnetic Resonance (1969). 1992 Jun;98(2):428–35.

9.	Neri D, Szyperski T, Otting G, Senn H, Wuethrich K. Stereospecific nuclear magnetic resonance assignments of the methyl groups of valine and leucine in the DNA-binding domain of the 434 repressor by biosynthetically directed fractional carbon-13 labeling. Biochemistry. 1989 Sep 19;28(19):7510–6.

10.	Hansen DF, Vallurupalli P, Kay LE. An Improved 15N Relaxation Dispersion Experiment for the Measurement of Millisecond Time-Scale Dynamics in Proteins. J Phys Chem B. 2008 May 1;112(19):5898–904.

11.	Ishima R, Torchia DA. Extending the range of amide proton relaxation dispersion experiments in proteins using a constant-time relaxation-compensated CPMG approach. J Biomol NMR. 2003;25(3):243–8.

12.	Yuwen T, Kay LE. Revisiting 1HN CPMG relaxation dispersion experiments: a simple modification can eliminate large artifacts. J Biomol NMR. 2019 Nov 23;73(10–11):641–50.

13.	Korzhnev DM, Kloiber K, Kanelis V, Tugarinov V, Kay LE. Probing Slow Dynamics in High Molecular Weight Proteins by Methyl-TROSY NMR Spectroscopy:  Application to a 723-Residue Enzyme. J Am Chem Soc. 2004 Mar 1;126(12):3964–73.

14.	Vallurupalli P, Bouvignies G, Kay LE. Studying “Invisible” Excited Protein States in Slow Exchange with a Major State Conformation. J Am Chem Soc. 2012 May 16;134(19):8148–61.

15.	Yuwen T, Sekhar A, Kay LE. Separating Dipolar and Chemical Exchange Magnetization Transfer Processes in 1H‐CEST. Angewandte Chemie International Edition. 2017 May 22;56(22):6122–5.

16.	Bouvignies G, Kay LE. A 2D 13C-CEST experiment for studying slowly exchanging protein systems using methyl probes: an application to protein folding. J Biomol NMR. 2012 Aug 12;53(4):303–10.

17.	Lundström P, Hansen DF, Kay LE. Measurement of carbonyl chemical shifts of excited protein states by relaxation dispersion NMR spectroscopy: comparison between uniformly and selectively 13C labeled samples. J Biomol NMR. 2008 Sep 2;42(1):35–47.

18.	Vallurupalli P, Kay LE. Probing Slow Chemical Exchange at Carbonyl Sites in Proteins by Chemical Exchange Saturation Transfer NMR Spectroscopy. Angewandte Chemie International Edition. 2013 Apr 8;52(15):4156–9.

19.	Long D, Sekhar A, Kay LE. Triple resonance-based 13Cα and 13Cβ CEST experiments for studies of ms timescale dynamics in proteins. J Biomol NMR. 2014 Dec 28;60(4):203–8.

20.	Bouvignies G, Vallurupalli P, Kay LE. Visualizing Side Chains of Invisible Protein Conformers by Solution NMR. J Mol Biol. 2014 Feb;426(3):763–74.

21.	Bolik-Coulon N, Hansen DF, Kay LE. Optimizing frequency sampling in CEST experiments. J Biomol NMR. 2022 Dec 4;76(5–6):167–83.

22.	Yuwen T, Kay LE. A new class of CEST experiment based on selecting different magnetization components at the start and end of the CEST relaxation element: an application to 1H CEST. J Biomol NMR. 2018 Feb 19;70(2):93–102.

23.	Hyberts SG, Takeuchi K, Wagner G. Poisson-Gap Sampling and Forward Maximum Entropy Reconstruction for Enhancing the Resolution and Sensitivity of Protein NMR Data. J Am Chem Soc. 2010 Feb 24;132(7):2145–7.

24.	Geen H, Freeman R. Band-selective radiofrequency pulses. Journal of Magnetic Resonance (1969). 1991 Jun;93(1):93–141.

25.	Massefski W, Redfield AG. Elimination of multiple-step spin diffusion effects in two-dimensional NOE spectroscopy of nucleic acids. Journal of Magnetic Resonance (1969). 1988 Jun;78(1):150–5.

26.	Bonin JP, Aramini JM, Kay LE. Structural Plasticity as a Driver of the Maturation of Pro-Interleukin-18. J Am Chem Soc. 2024 Nov 6;146(44):30281–93.

27.	Jing B, Berger B, Jaakkola T. AlphaFold Meets Flow Matching for Generating Protein Ensembles. ArXiv. 2024 Sep 2;

28.	Bonin JP, Aramini JM, Dong Y, Wu H, Kay LE. AlphaFold2 as a replacement for solution NMR structure determination of small proteins: Not so fast! Journal of Magnetic Resonance. 2024 Jul;364:107725.

29.	Mirdita M, Schütze K, Moriwaki Y, Heo L, Ovchinnikov S, Steinegger M. ColabFold: making protein folding accessible to all. Nat Methods. 2022 Jun 30;19(6):679–82.

30.	Teixeira JMC, Liu ZH, Namini A, Li J, Vernon RM, Krzeminski M, et al. IDPConformerGenerator: A Flexible Software Suite for Sampling the Conformational Space of Disordered Protein States. J Phys Chem A. 2022 Sep 8;126(35):5985–6003.

31.	Shen Y, Delaglio F, Cornilescu G, Bax A. TALOS+: a hybrid method for predicting protein backbone torsion angles from NMR chemical shifts. J Biomol NMR. 2009 Aug;44(4):213–23.

32.	Vallurupalli P, Bouvignies G, Kay LE. A Computational Study of the Effects of 13C– 13C Scalar Couplings on 13C CEST NMR Spectra: Towards Studies on a Uniformly 13C‐Labeled Protein. ChemBioChem. 2013 Sep 23;14(14):1709–13.
